# Challenges and opportunities in detecting leaf water and carotenoid content across biomes from satellite multispectral indices

**DOI:** 10.1101/2025.11.01.686024

**Authors:** Martha G. Montes-Bojórquez, Carlos A. Robles-Zazueta, Clara Tinoco-Ojanguren, Brian J. Enquist, Amy E. Frazier, Brian S. Maitner, Gabriel Massaine Moulatlet, José Raúl Romo-León, Lei Song, César Hinojo-Hinojo

## Abstract

Climate change is causing vegetation stress across the globe, increasing the need for reliable indicators to monitor plant health. Leaf water and carotenoid content, and the chlorophyll/carotenoid ratio, are established proxies for environmental stress that can be detected by remote sensing. Here, we evaluated the sensitivity of 11 multispectral vegetation indices (VIs) designed to monitor these three stress-related leaf traits across a broad range of environmental and vegetation conditions. For this, we combined radiative transfer modeling with cross-biome field and satellite observations from Sentinel-2, Landsat 8, and MODIS from the National Ecological Observatory Network (NEON), spanning in most major terrestrial ecosystems. Our model-based analysis showed that VIs have a low to moderate sensitivity to their target traits, ranging from water indices with 66% of their variability explained by leaf water content, to carotenoid indices with 27% variability explained by leaf carotenoid content. Surprisingly, our field-based analyses revealed minimal to no sensitivity to leaf water and carotenoid content and chlorophyll/carotenoid ratio across all VIs. In contrast, we showed that leaf area index was the dominant driver of all studied VIs, accounting for 54-74 % of their variability in the field-based analysis. Lastly, we detected that VIś sensitivity to atmospheric conditions and field sampling issues contribute to their low performance in validating ground truth observations. These findings show that improvements in the VIs formulation and field sampling strategies are needed to increase the reliability of vegetation stress monitoring from multispectral satellites and support a generalized use of VIs across ecosystems.

****Highlights:** 3-5 bullet points, 85 characters:**

● Sensitivity of water and carotenoid multispectral indices was evaluated
● Analysis based on cross-biome field data and radiative transfer models
● Field data showed indices had minimal sensitivity to leaf water and carotenoid
● Leaf area index explained most cross-biome variation in water and carotenoid indices
● We propose strategies to improve stress-related index formulation and validation

## 1. INTRODUCTION

The effects of climate change, such as increasing frequency and intensity of heatwaves, frosts, or prolonged droughts, are having significant impacts on vegetation globally. Such events have led to widespread plant mortality, reduction in plant productivity, changes in species-area distributions, and changes in community species composition (Singh et al. 2021; Hammond et al. 2022). Climatic shifts trigger various physiological changes as part of their response mechanisms to thermal and water stress (Zandalinas & Mittler, 2022). These responses, which reflect trait plasticity, include reduced leaf area, root system expansion, and alterations in leaf pigment concentrations (Fahad et al. 2017; Yang et al. 2021). Despite advances in plant stress research, traditional methods for directly assessing morphological and physiological changes remain time-consuming, costly, and often invasive, making them impractical for large-scale or long-term monitoring. Thus, plant and remote sensing scientists have been on a quest for reliable proxies to monitor plant stress status and anticipate the cascading impacts of climate change on vegetation (Gamon et al. 2016, Boving et al. 2025).

Two traits that change rapidly under stress are leaf water content (LWC) and foliar pigments, especially carotenoids. LWC is a key trait that reflects a plant’s water status and provides valuable information on drought response across developmental stages. It is associated with leaf water potential, photosynthesis, stomatal conductance, and transpiration (Wang et al. 2022; Yasir et al. 2024). Carotenoids are also highly sensitive to environmental conditions and tend to increase when the canopy is subject to abiotic stresses due to their photoprotective and antioxidant role (Gamon et al. 2016, Wong et al. 2019). More specifically, carotenoids contribute to photoprotection by enhancing light-harvesting and by dissipating excess absorbed energy, and prevent oxidative stress in the photosynthetic machinery by neutralizing stress-related reactive oxygen species (Dall’Osto et al. 2014; Simkin et al. 2022; Murchie & Lawson, 2013; Gamon et al. 2016; Sun et al. 2022). The water molecules absorb radiation in the near and shortwave infrared (1350-2500 nm), while carotenoids absorb in the visible (500 to 550 nm) regions commonly captured in hyperspectral and multispectral space-borne imagery (Jacquemoud et al. 2009, Feret et al. 2008). Such optical properties make these traits good proxies of plant health status and strong candidates for remote sensing monitoring at large scales and across time.

Vegetation indices (VIs) are mathematical formulations that combine reflectance spectral bands from multi- or hyperspectral sensors, designed to highlight information on key biophysical properties (Xue & Su, 2017; Sims & Gamon, 2002; Xue & Su, 2017). VIs have become a fundamental tool for monitoring vegetation stress due to their usefulness, accessibility, low computational demand, and ability to provide low-cost information across broad spatial and temporal scales (Verrelst et al. 2015). Several vegetation indices have been developed to be sensitive to water and carotenoid content in the vegetation, such as the Normalized Difference Water Index (NDWI), Carotenoid Reflectance Index (CRI_550_), and the Chlorophyll/Carotenoid Index (CCI) (Gitelson et al. 2002; Ji et al 2011; Gamon et al 2016; Yin et al. 2022). The ability of multispectral water indices to detect leaf water content in vegetation has been ground-validated in local settings, with poor to moderate performance, including single canopies with varied leaf and canopy properties (Sims & Gamon 2003, Zarco-Tejada et al 2003). Other studies have highlighted that water indices could have an improved ability to detect variation in the leaf area index (LAI), compared to other commonly used indices like NDVI (see Sims & Gamon 2003). Previous sensitivity analyses of radiative transfer models further confirm that several water indices are mainly sensitive to both leaf water content and LAI (Morcillo-Pallarés et al. 2019; Xiao et al 2014). Similarly, multispectral carotenoid indices have been ground-validated on a few local sites (e.g., Gamon et al., 2016 for CCI), and to the best of our knowledge, they have not been thoroughly assessed with sensitivity analyses of radiative transfer models. More importantly, the ability of multispectral water and carotenoid indices to capture leaf water and carotenoid content has not been broadly tested with ground data across a wide range of biomes and environmental conditions. Despite the lack of broad and thorough ground validation, the generalized use of multispectral water and carotenoid indices has increased as indicators of water stress, and the physiological status of the vegetation (Yin et al. 2022; Wang et al 2023, Hais et al 2019, Xu et al 2025), highlighting the need for a rigorous broad-scale assessment that supports its widespread use.

The generalized use of VIs without broad-scale validation, while a common practice in remote sensing science, can lead at least to attribution errors. Such errors can arise because the spectral response of vegetation is also influenced by a range of leaf and canopy biochemical and structural traits, soil characteristics, atmospheric conditions, sun and sensor geometries (Asner 1998, Ollinger 2011). These factors may also vary depending on the ecosystem (Asner 1998), which can result in artifacts or errors in the interpretation of vegetation stress and in inaccurate estimations of foliar traits when indices are applied in variable contexts. Understanding these limitations is essential for improving the use of VIs under diverse environmental conditions.

Here, we evaluate the ability of existing multispectral water and carotenoid VIs estimated from satellite imagery to detect the leaf water and carotenoid content. To achieve this, we assessed the performance of these VIs using radiative transfer modeling (RTM) and ground truth data from sites spanning most major terrestrial ecosystem types. We hypothesize that the sensitivity and performance of these VIs vary depending on environmental and structural conditions, which limits their general applicability.

## 2. MATERIALS AND METHODS

### 2.1. Selection of multispectral indices

We evaluated 11 VIs sensitive to carotenoids and water content based on the literature and the Index Database (IDB: www.indexdatabase.de). Our analysis focused on VIs designed for multispectral sensors compatible with data from MODIS, Sentinel-2, and Landsat 8. The only exceptions are the CRI and Car indices, which were originally developed for hyperspectral data at leaf scale but use spectral regions that fall within Sentinel-2 sensor bands, and given their good performance at leaf level and in our pilot model-based explorations, we considered it worth including them. The 11 VIs were evaluated using global sensitivity analysis, which quantifies how variation in model input parameters affects simulated reflectance, and validated using ground truth data from a wide range of terrestrial ecosystems, as described below (Table 1).

**Table 1.** Summary of the 11 vegetation indices evaluated in this study, including their formulas, corresponding sensors, and references. Indices are grouped according to the biochemical/physiological trait they are primarily associated with (leaf water content, leaf carotenoid content, and leaf chlorophyll/carotenoid ratio). In the index formulas, *R* denotes reflectance at a specific wavelength, while b refers to reflectance in a spectral band. For sensors such as Sentinel-2, where certain wavelengths are not directly available, the closest central band was used.

| Biochemical/<br>Physiological<br>trait | Index | Formula | Sensor | Reference |
| --- | --- | --- | --- | --- |
| Leaf water<br>content | Normalized difference water index (NDWI) | $\frac{(R860 - R1240)}{(R860 + R1240)}$ | MODIS | Zarco-Tejada et al. 2003 |
| | Simple ratio water index (SRWI) | $\frac{R860}{R1240}$ | MODIS | Zarco-Tejada et al. 2003 |
| | Normalized difference moisture index (NDMI) | $\frac{(R830 - R1650)}{(R830 + R1650)}$ | Landsat 8 | Suming & Sader, 2005 |
| | Tassalled Cap - wetness (TCW) | $(0.1509 * b2) + (0.1973 * b3)$ | Landsat 8 | IDB; Suming & Sader, 2005 |
| Leaf carotenoids<br>content | Carotenoid reflectance index (CRI<sub>550</sub>) | $\left(\frac{1}{R510}\right) - \left(\frac{1}{R550}\right)$ | Sentinel-2 | Gitelson et al. 2002 |
| | Carotenoid reflectance index (CRI<sub>700</sub>) | $\left(\frac{1}{R510}\right) - \left(\frac{1}{R700}\right)$ | Sentinel-2 | Gitelson et al. 2002 |
| | Red edge carotenoid index (CAR<sub>red-edge</sub>) | $\left[\left(\frac{1}{R510}\right) - \left(\frac{1}{R700}\right)\right] * R770$ | Sentinel-2 | Gitelson et al. 2006 |
| | Green carotenoid index (CAR<sub>green</sub>) | $\left[\left(\frac{1}{R510}\right) - \left(\frac{1}{R550}\right)\right] * R770$ | Sentinel-2 | Gitelson et al. 2006 |
| Leaf<br>chlorophyll/carot<br>enoid ratio | Chlorophyll/carotenoid index (CCI) | $\frac{(R531 - R645)}{(R531 + R645)}$ | MODIS | Gamon et al. 2016 |
| | Structure insensitive pigment index (SIPI) | $\frac{(R800 - R445)}{(R800 + R680)}$ | Sentinel-2 | Peñuelas et al. 1995 |
| | Green-red vegetation index (GRVI) | $\frac{(R555 - R645)}{(R555 + R645)}$ | Landsat 8 | Yin et al. 2022 |

### 2.2. Model-based evaluation of multispectral indices

We performed a variance-based global sensitivity analysis of the radiative transfer models PROSPECT-D and 4SAIL for each of the selected multispectral indices from Table 1. The PROSPECT-D model is a well-calibrated biophysical representation of the radiation absorbance, reflectance, and transmittance of a single leaf given its values for various biochemical and structural traits, including its water and carotenoid content (Féret et al. 2017). The 4SAIL model predicts surface reflectance through a simplified representation of the canopy, incorporating the effect of leaf reflectance, canopy structure (LAI and mean leaf inclination angle), soil reflectance, sun and sensor geometries, among others (Jacquemoud et al. 2009). Altogether, the PROSPECT-D and 4SAIL model parameters represent the major factors controlling surface reflectance through well-established optical properties (Féret et al. 2008; Jacquemoud et al. 2009). We coupled both models through the global sensitivity analysis. Briefly, a variance-based global sensitivity analysis varies a number of times all model input parameter values together across a given set of ranges and values distribution, and statistically assesses how much variability of the model output is explained by each input parameter and their interactions, i.e, estimating the outputs’ sensitivity to varying input parameters. Here, the output was the VI values estimated from the modeled surface reflectance at the canopy scale, and we assessed the VIs’ sensitivity to variation in PROSPECT-D and 4SAIL parameters. The ranges at which each parameter was allowed to vary and their rationale are shown in Table 2. The leaf and canopy ranges are based on the mean value of the canopy found across the field sites included in this study (Table 2, Supplementary Table S1). A total of 2,000 parameter value combinations were generated using the Sobol sampling distribution (Saltelli et al. 2010), which assigns values for each parameter in a way that all value combinations altogether uniformly cover the whole parameter space. Each parameter value combination represents a canopy with a given set of trait values and environmental and sun/sensor geometry conditions. For example, a conifer forest would be a combination of mid to high leaf area index, high leaf mass per area, and mid to high leaf chlorophyll, carotenoid, and water contents. All parameter combinations and this analysis enabled us to evaluate the sensitivity of VIs to these traits across a wide range of environmental conditions and ecosystems, covering variations in the structural and biochemical properties of the canopy and leaves in a way that represents the extent of variation found across the major terrestrial vegetation types. This analysis was conducted using the ARTMO toolbox and its Global Sensitivity Analysis module (Verrelst et al. 2015).

**Table 2.** PROSAIL (PROSPECT-D + 4SAIL) parameter ranges used in the global sensitivity analysis. Values correspond to biochemical, structural, and observational traits defined based on the abundance-weighted average values estimated from the cross-biome NEON field data.

| Parameter | Min | Max | Rationale |
| --- | --- | --- | --- |
| Leaf chlorophyll content ($\mu\text{g cm}^{-2}$) | 11 | 85 | Range found at the study sites* |
| Leaf carotenoid content ($\mu\text{g cm}^{-2}$) | 3 | 30 | Range found at the study sites* |
| Leaf structure (unitless) | 1 | 4 | Ranges from Jacquemoud and Baret (1990) |
| Leaf mass per area ($\text{g m}^{-2}$) | 48 | 552 | Range found at the study sites* |
| Leaf water content ($\text{g m}^{-2}$) | 10 | 771 | Range found at the study sites* |
| Leaf area index (m<sup>2</sup> m<sup>-2</sup>) | 0.2 | 7 | Exploration global and study sites data** |
| Mean leaf angle (°) | 25 | 63 | Global multi-biome data synthesis *** |
| Hotspot | 0.0001 | 0.01 | Estimated variation across terrestrial ecosystems**** |
| Soil reflectance | 0 | 1 | All possible range |
| Diffuse:direct radiation proportion | 0 | 1 | All possible range |
| Solar zenith angle (°) | 21 | 76 | Exploration of Landsat, MODIS, and Sentinel-2 global data |
| Sensor view angle (°) | 0 | 0 | Multispectral sensors have a near-nadir view or estimated at nadir |
| Relative azimuth angle (°) | 0 | 180 | All possible range |

\*Based on variation found at plot data from our study sites, estimated as the community weighted mean (weighted by species abundance). See section Field-based evaluation of multispectral indices
\*\*Minimal value was set to 0.2 to avoid using 0 (no leaves). The maximum value was set to the approximate maximum value found at a global exploration of leaf area index MODIS MOD15A2H product and field data at study sites.
\*\*\*Uses the 10 and 90% quantiles of a multi-biome global data synthesis from Hinojo-Hinojo and Goulden (2020b), which closely aligns with other independent global data from Li and Fang (2025).
\*\*\*\*According to the hypothesized cross-biome variation of the ratio of leaf width to canopy height (Hinojo and Goulden 2020)

Once the traits to which the VIs are sensitive were identified, an exploratory analysis using the PROSAIL modeled data (using the same settings as the global sensitivity analysis) was done to better describe how variation in the more important traits actually influences the VIs. To better describe the trait-VI relationship, we fit the modeled data with linear, power, and exponential regressions. Then, LAI was grouped into three categories: (1) full range, from 0.2 to 7; (2) intermediate, from 2 to 7, which excludes the lowest LAI values; and (3) high, from 4 to 7, which includes only the upper LAI range. These ranges represent increasing canopy density, from sparse to dense vegetation cover. This exploratory analysis was conducted with the “hsdar” package (Lehnert et al. 2019) for R software.

### 2.3. Field-based evaluation of multispectral indices

Radiative transfer modeling is increasingly used to confirm VIs’ sensitivity to their target vegetation properties under a broad range of conditions (Morcillo-Pallarez et al., 2019; Yan et al., 2025). However, a thorough evaluation of such sensitivity with field data from a broad range of terrestrial biomes is still lacking in several multispectral VIs, including water and carotenoid indices. To address this issue, we integrated satellite information with ground truth data from diverse terrestrial ecosystems across the US, where the major traits known to influence surface reflectance have been measured with standardized protocols. This data will be used to further complement the model-based global sensitivity analysis to assess if the VIs are actually sensitive to leaf water or carotenoid content measured in the field.

We obtained field data from the National Ecological Observatory Network (NEON, https://www.neonscience.org/), a major ecological monitoring network encompassing the continental United States, Hawaii, and Puerto Rico. We extracted key vegetation traits that influence surface reflectance, including leaf carotenoid, water, and chlorophyll content, and leaf mass per area from the plant foliar traits product (DP1.10026.001); species abundance from the vegetation structure product (DP1.10098.001); and leaf area index (LAI) data from the digital hemispheric photos of plot vegetation product (DP1.10017.001) as processed by the Ground-Based Observations for Validation of Copernicus Global Land Products (https://gbov.land.copernicus.eu/). In addition, LAI estimates were derived from the MODIS MCD15A3H product using Google Earth Engine. We collected these measurements from 193 plots across 39 monitoring sites, representing major terrestrial ecosystems including tropical and temperate forests, deciduous, mixed, and coniferous forests, as well as shrublands, tundra, grasslands, and crop fields (see Supplementary Table S1).

For each site, we calculated the vegetation’s average leaf trait values by weighting trait data by the species’ relative basal area, as a surrogate for species biomass contribution. We used only the data for the plots that NEON tag as “tower” plots, which are the most frequently and thoroughly sampled and have more supporting information available (e.g., phenocam photos of over- and understory to confirm vegetation physiognomy and structure). We discarded all plots tagged as “distributed” plots. For plots dominated by grass or herbaceous vegetation, the NEON sampling already provides an abundance-weighted mean for each sampled trait per plot, so no further processing was needed (Weintraub-Leff, 2024). We discarded sites and plots where both woody and herbaceous plants were dominant components in the landscape (e.g., open woodlands with a significant understory layer), as we lacked a consistent way to properly weight the abundance of both components (Hinojo-Hinojo et al. 2024). We only kept those plots where we had trait data for at least 80% of the species stem area, as a criterion to ensure vegetation was well represented (Pérez-Harguindeguy et al. 2013). We estimated the statistical distribution of each trait by combining the trait and species basal area data for each plot using a state of the art method based on a bootstrap sampling that explicitly incorporates intraspecific variation (Maitner et al., 2023). We estimated the mean trait value of the vegetation from such distributions. These estimations were performed with the “traitstrap” R package (v. 0.1.0, Maitner et al. 2023).

Surface reflectance data were obtained from Landsat 8, MODIS, and Sentinel-2 satellite missions via Google Earth Engine. We extracted the values corresponding to the single pixel located at the coordinates of the NEON plots within a time window of 32 days centered around the mid date of the trait sampling. Such a window ensured that data from at least 3 Landsat-8 images were included. Landsat 8 and Sentinel-2 pixel size roughly compares to the plot size at most sites with woody vegetation (NEON plots are approximately 40 × 40 meters). In contrast, while it can be significantly larger for the grassland or herbaceous plots, where we assumed the mean trait value of such plots would approximate the mean value at the whole pixel. We extracted the values of the pixel containing the location of each NEON plot within a two-week time window centered on the mid date of trait sampling. For MODIS data, we used images from the MCD19A1 product (version 061), which provides daily surface reflectance data for MODIS land and ocean bands corrected for atmospheric conditions and bidirectional reflectance factor. Given that the pixel size of such MODIS products (500 to 1000 m, depending on the band) is considerably larger than the plot size, we extracted reflectance data for the single pixel containing the larger majority of each site’s field plots, so that pixel is sampled by several plots. To ensure data quality, all images were filtered to remove pixels affected by clouds, cloud shadows, or snow, using the quality assurance bands provided by each satellite product. These reflectance values from different spectral bands were then used to calculate the VIs. Lastly, we obtained the average value of each VI for each plot (or site for MODIS) for the 32-day time window around the sampling date, which were then merged into a single database containing the field and satellite VIs data for each plot/site.

### 2.4. Data Analysis

For the modeled data, we analyzed the ability of the indices to estimate water and carotenoid content using regression models. Power functions were used for most relationships, as they provided a better fit to the data than linear models, while an exponential model was used for CCI. We only show the model type that best captures the observed trend in the data. Then, for the field-based analysis, we explored the factors influencing these indices through multiple regression analyses, using satellite-derived index values as response variables and field trait estimates (e.g., leaf water, chlorophyll and carotenoid content, LAI) as predictors. Finally, we performed variance partitioning analyses based on ground truth and satellite data, in order to assess how much of the total variation in each VI could be attributed to leaf water or pigments content, or other key traits of the leaves and the canopy, such as LAI or LMA. For each VI, we used the index value as the response variable and a combination of key vegetation traits (e.g., water content, LAI, and LMA for NDWI) as explanatory variables. The variance partitioning analyses were performed using the R package “vegan” (v. 2.6-10, Oksanen et al. 2025). All analyses were conducted in R (R Core Team 2025).

## 3. RESULTS

### 3.1. Modeling multispectral indices sensitivities

Our global sensitivity analysis using the PROSAIL model revealed that, over a broad range of vegetation conditions, the target traits (either leaf water and carotenoid content, chlorophyll/carotenoid ratio) explained less than half of the total variability of the VIs (Figure 1). Water content explained over 40% of the water index variability. Of these four, the Tasseled Cap - Wetness (TCW) index was the one with the largest sensitivity to variation in leaf water content, explaining 66.14% of the VI variability, and it was also the water VI least affected by LAI. Leaf carotenoid content explains only about 1/4th of the variation of the best performing carotenoid index (CRI_550_). On the other hand, Car_rededge_ was the carotenoid index less sensitive to variation in carotenoid content, with only 18.97% (Figure 1). Of the three VIs analyzed for the chlorophyll/carotenoid ratio (CCI, SIPI, and GRVI), only CCI was sensitive to both pigments, which explains nearly 40% and 25% of this index variability, respectively. SIPI and GRVI were largely insensitive to carotenoid content (Figure 1) but had a low to moderate sensitivity to chlorophyll. The sensitivity analysis also revealed that VIs have a substantial sensitivity to LAI, while also showing a lower sensitivity to LMA, leaf structure, and soil reflectance (Figure 1).

**Figure 1.**
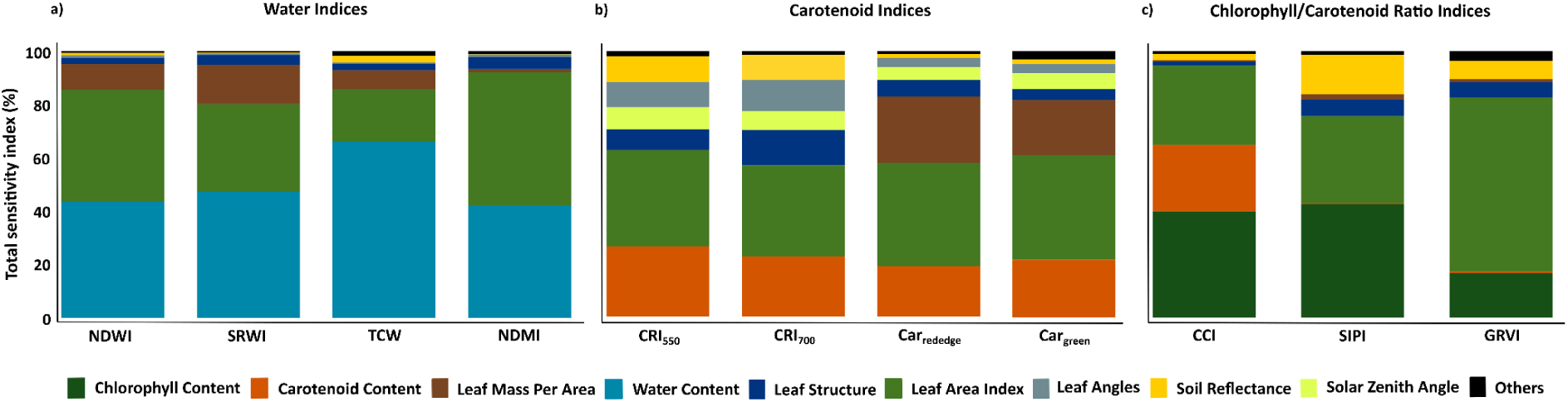
Percent of variance (0-100%) of multispectral indices explained by key leaf and canopy traits, environmental conditions, and sun/sensor geometry according to a global sensitivity analysis of the PROSAIL model. Each bar represents a multispectral VI (VIs grouped by target trait: leaf water content, leaf carotenoids, and leaf chlorophyll/carotenoid ratio). The colors indicate the relative contribution of different biophysical parameters to the total variation of each index, i.e., the index sensitivity to a given parameter. A value close to 100% would mean that most variation in the index is explained by a given parameter, while lower values (e.g., 40%) indicate limited explanatory capacity and higher influence from other parameters/factors. Accordingly, higher bars denote greater sensitivity of the index to the target trait, whereas smaller proportions reflect a lower capacity to capture variation in that trait.

We found that the VIs’ sensitivity to leaf water and carotenoid content, and chlorophyll/carotenoid ratio increased substantially, and the sensitivity to LAI decreased, at LAI > 2, a level above which VIs saturate to LAI (Figure 2, and Figures S1-S3). At LAI> 2, the relationship of VIs and their target trait becomes stronger and practically linear, except for CCI whose relationship remains exponential but substantially stronger (Figure 2b-c, e-f, g-i, Figures S1-S3). The indices that showed the best model fits were: for water content, the Normalized Difference Water Index (NDWI, R² = 0.85), for carotenoid content, the Carotenoid Reflectance Index (CRI_550_, R² = 0.50), and for the chlorophyll/carotenoid ratio, the Chlorophyll Carotenoid Index (CCI, R² = 0.79).

**Figure 2.**
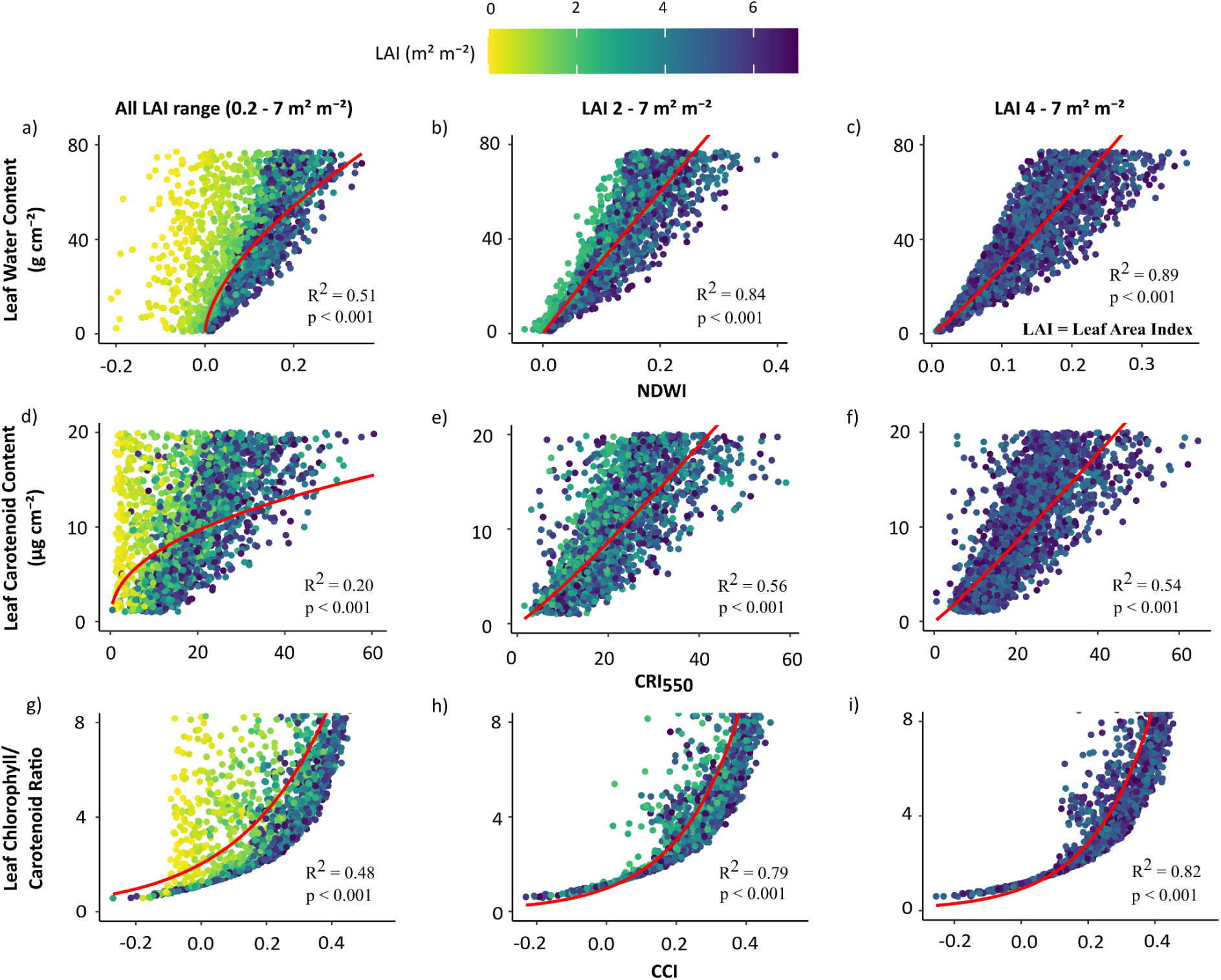
Influence of leaf area index (LAI) on the relationships between NDWI, CRI_550_, and CCI and their target traits (leaf water content, leaf carotenoids content, and leaf chlorophyll/carotenoid ratio), according to the PROSAIL model. Each point represents a single combination of parameter values generated with the PROSAIL model, i.e., a canopy with particular values of leaf and canopy traits, and environmental and sun/sensor geometry conditions. The color gradient (yellow to dark blue) indicates increasing LAI values, from sparse to dense canopies. Columns correspond to three LAI ranges: (a, d, g) full range (0.2–7), (b, e, h) medium range (2–7, excluding low LAI), and (c, f, i) high range (4–7). We based this categorization on the observation that the strongest dispersion occurred at low LAI levels, while higher values were more consistent.

### 3.2. Cross-biome assessment of multispectral indices with satellite and field data

Our variance partitioning analysis revealed that the studied VIs had minimal to null sensitivity to the leaf water and carotenoid content and the chlorophyll/carotenoid ratio in the field. These traits explained less than 8 % of the VIs’ variation across all plots and sites from a broad range of ecosystems (Figure 3 and Figures S4-S6), CRI_550_ and CCI are the only one showing a statistically significant relationship with its target trait, although very weakly (Figure 4 and Figures S7-S9). Such low sensitivity is not clearly related to the presence of sparse vegetation (low LAI, Figure 4a-c and Figures S7-9), as our modeling analysis suggested (Figure 2a, c, and g). In contrast, LAI consistently emerged as the primary driver of the observed variation of the magnitude of the studied VIs (57-74 % variation, Figure 3), showing a positive, almost linear relationship (Figure 4d-f).

**Figure 3.**
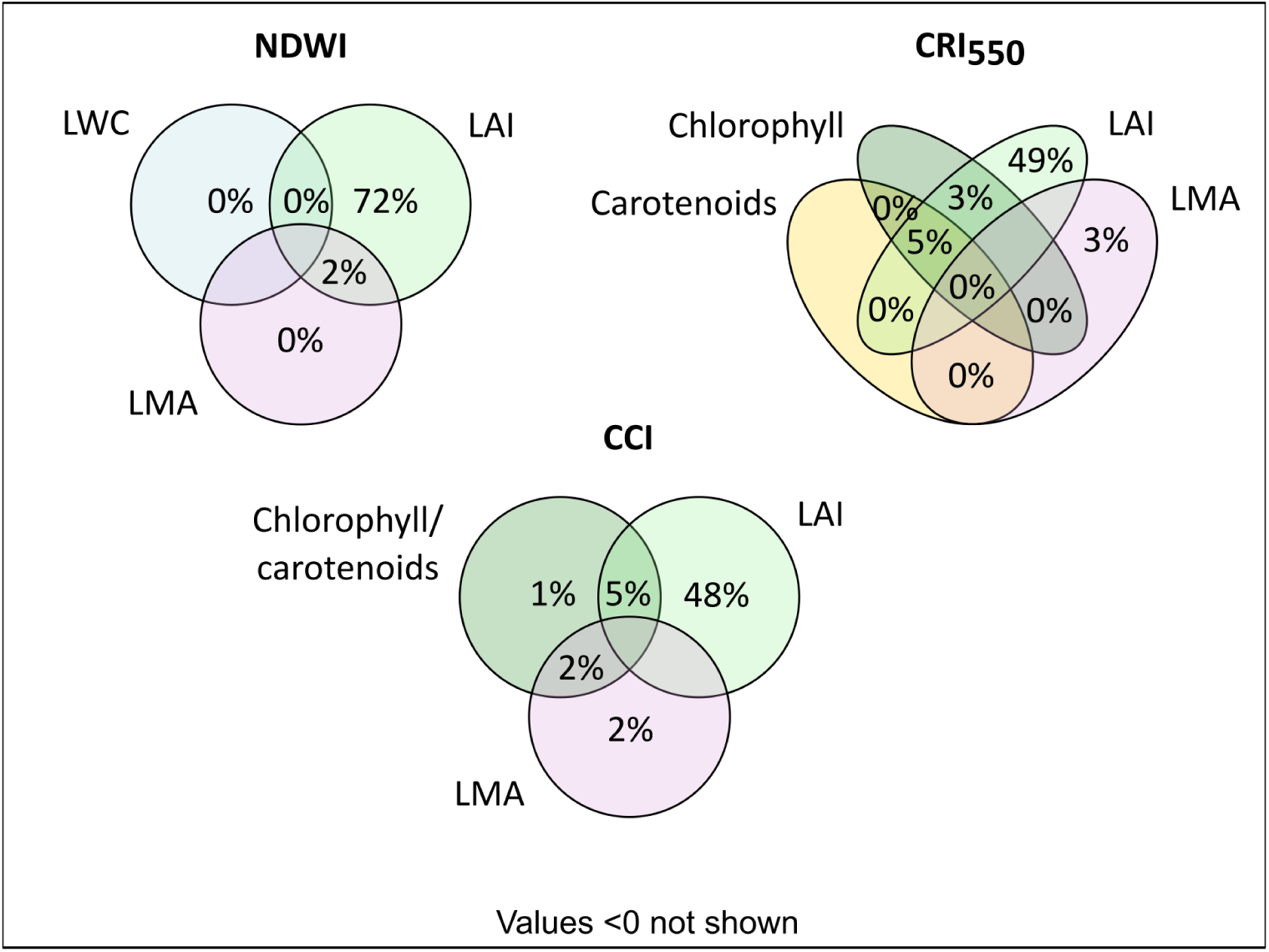
Venn diagrams of the proportion of observed variation in three multispectral indices—NDWI, CRI_550_, and CCI—explained by key leaf and canopy traits, according to variance partitioning analysis of ground-truth and satellite data. Each circle represents a predictor variable included in the models: for NDWI, leaf water content (LWC), leaf area index (LAI), and leaf mass per area (LMA); for CRI_550_, carotenoids, chlorophyll, LAI, and LMA; and for CCI, chlorophyll, carotenoids, and LAI. Numbers indicate the percentage of variance in each index uniquely or jointly explained by these traits. Non-overlapping areas denote independent contributions, while overlaps indicate shared explanatory effects among variables.

**Figure 4.**
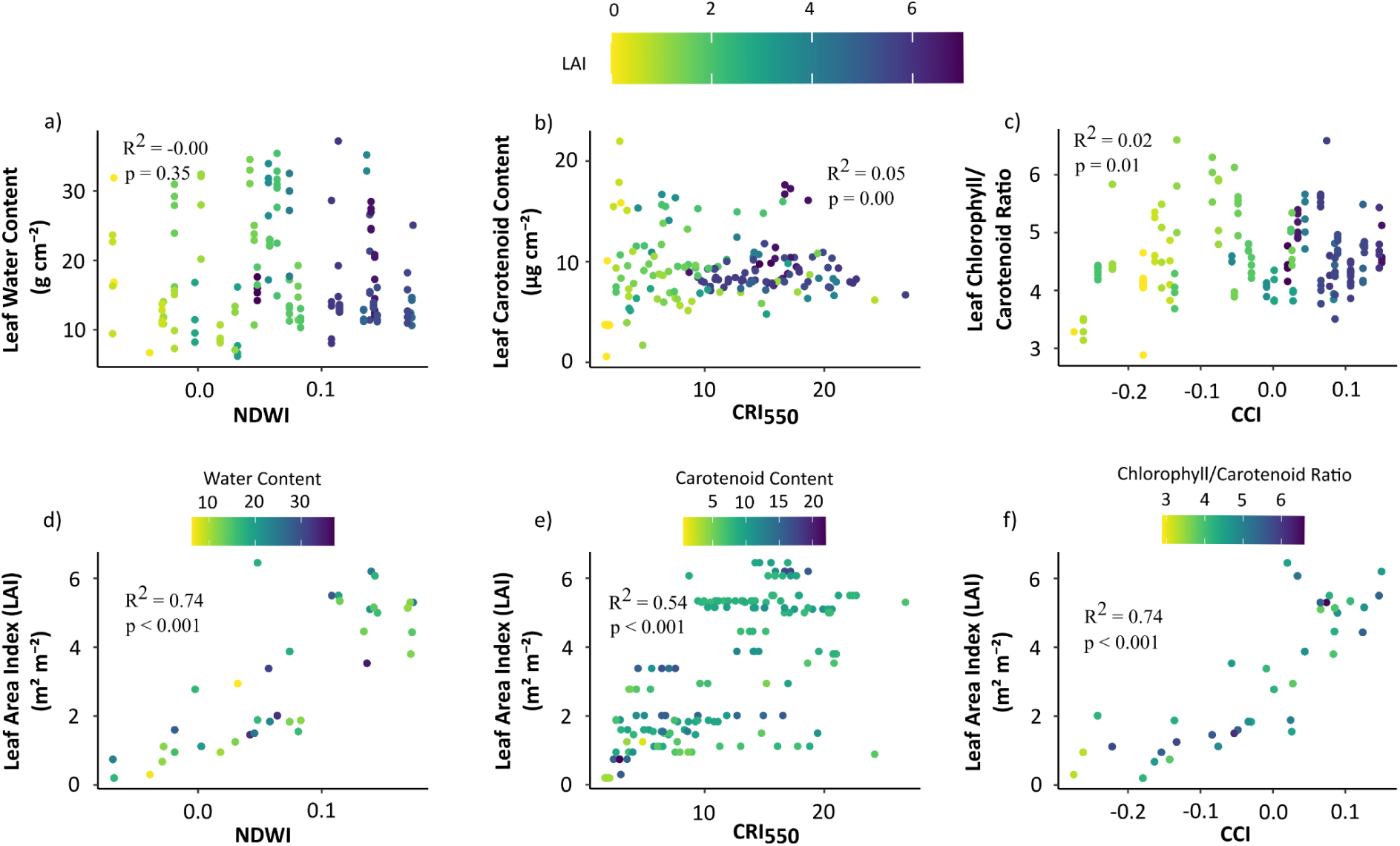
Relationships between three multispectral indices—NDWI, CRI_550_, and CCI—and their corresponding leaf traits (a–c) and leaf area index (LAI) (d–f), based on ground-truth and satellite data. Each point represents a field–satellite-paired observation. Panels (a–c) show weak or no relationships between indices and their target biochemical traits (leaf water content, leaf carotenoids content, and leaf chlorophyll/carotenoid ratio), whereas panels (d–f) reveal strong relationships with LAI, indicating that index variation is mainly driven by canopy structure rather than leaf pigment or water content.

### 3.3. Serendipitous factors influencing the indices vs leaf trait relationships

We detected a sampling bias in some NEON open forest sites where the understory is abundant but its presence or importance is not obvious from the abundance or trait data, as the understory was not sampled at all. We detected this after careful exploration of sites’ phenocam and aerial imagery in NEON sites such as SOAP (Soaproot Saddle), DEJU (Delta Junction), YELL (Yellowstone National Park), RMNP (Rocky Mountains National Park), JERC (The Jones Center At Ichauway), LENO (Lenoir Landing), CLBJ (Lyndon B. Johnson National Grassland), and BONA (Caribou-Poker Creeks Research Watershed). We illustrate this with aerial photos of the YELL site (Figure 5a). Plots in this site show widely varying degrees of evergreen trees and herbaceous understory composition, but understory species were not sampled for leaf traits. We observed three patterns that are at least partly related to that sampling bias. First, plots at some sites show relatively consistent leaf water or carotenoid contents but more widely varying VI values, indicative of plots with similar tree species composition (and so similar sampled values of leaf contents) but with varying degrees of tree/understory abundance (understory variation not sampled but captured by VIs). This pattern is notorious at YELL site (Figure 5c) where leaf carotenoid content is about 6 µg cm^-2^ but CRI550 varies widely (from 7 to 15), but also present at some plots at RMNP and DEJU sites in Figure 5b and at most plots at the SOAP and in Figure 5c. Second, plots at some sites show fairly consistent VI values but widely varying leaf water or carotenoid content, indicative of sites where uniform understory may dominate the spectral VI signal (but was not sampled for leaf traits), but low-abundance tree species vary among plots (sampled, with varying leaf contents). This can be observed in most plots at the RMNP site (Figure 5b), where NDMI plot values are about 0.18 while leaf water content had more variation (∼20 to ∼ 33 g m^-2^), and also at YELL site. Third, most of the plots and sites where we identified abundant but unsampled understory are located at the upper or lower extreme of the scatterplot, a potential indicator of such biased sampling (Figure 5b-c).

**Figure 5.**
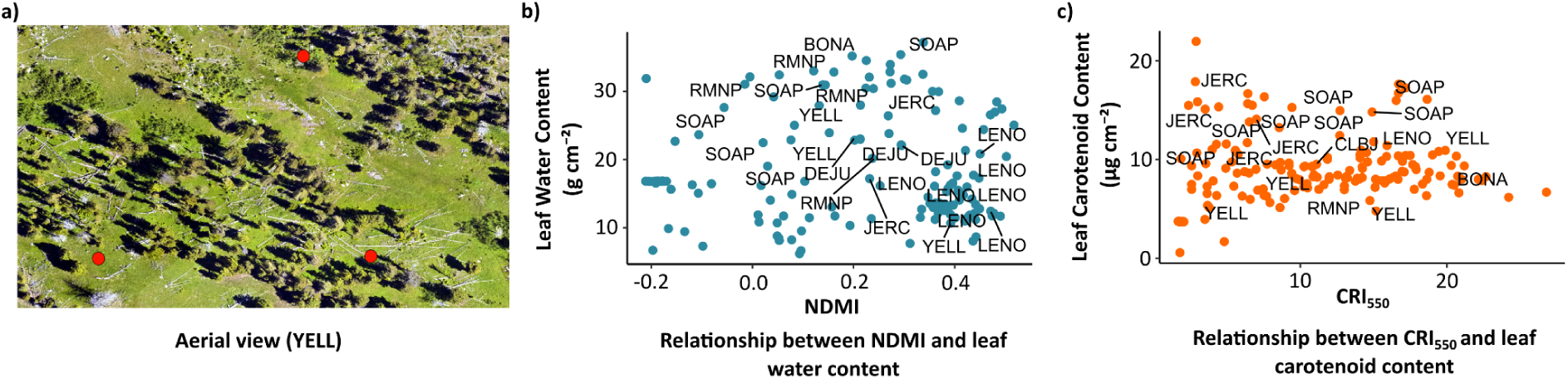
Example of incomplete and spatially heterogeneous sampling in open forests with understory. (a) Aerial view of YELL site showing plots (red circles) in locations with varying abundance of rich understory and trees. (b) Variation in leaf water content and NDMI, and (c) leaf carotenoid content and CRI_550_, across all field plots, highlighting the sites that have both trees and abundant understory present, where understory was not sampled for traits or abundance. The influence of such sampling bias (not sampling the abundant understory) can be seen in (b) and (c) as 1) plots within a site with similar leaf contents but widely different VI values, 2) plots within a site with similar VI values but widely different leaf content values, and 3) plots skewed to the high or low portion of the plots area..

At several sites, notorious reductions in VIs values within the peak growing season coincided with the presence of clouds or thin cloud layers that were not detected by the cloud removal algorithm (Figure 6c). We illustrate this with the JERC site, where CRI_550_ varied markedly, including a sharp decrease on May 23, 2024 (Figure 6a). The RGB satellite image from the same acquisition date reveals a whitish, opaque tone over the plot location, indicative of a thin cloud layer, while more defined clouds are visible in the surrounding areas (Figure 6c). This atmospheric interference, although subtle, alters the spectral signal, producing an artificial decrease in carotenoid index values of nearly half the range of CRI₅₅₀ found across all sites (Figure 6a, see full cross-site CRI_550_ range in Figure 4b and 4e). For comparison, we show that NDVI was much less sensitive than CRI₅₅₀ with much less pronounced fluctuations throughout the same time period (Figure 6b). These results suggest that CRI₅₅₀ can be highly sensitive to atmospheric disturbances, and that thin cloud layers, even when barely perceptible, may account for part of the mismatch between satellite VI data and our leaf trait field estimates.

**Figure 6.**
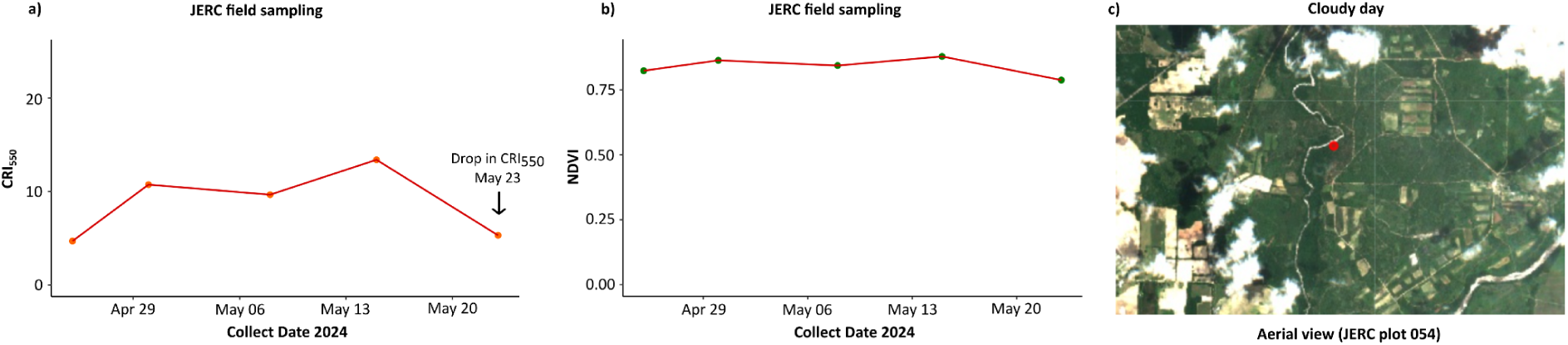
Example of the influence of atmospheric noise over CRI_550_ at a single plot at the JERC site. (a) CRI_550_ shows a sharp decrease on May 23. (b) NDVI remains stable during the same period. (c) Sentinel-2 RGB image of May 23, where a whitish opaque tone over the plot (red dot) suggests the presence of a very thin cloud cover, while thicker clouds are visible in the surrounding area. Overall, the figure shows that CRI550 is very sensitive to slight atmospheric noise, even more than NDVI.

In some NEON sites, the field sampling for leaf traits was completed within a few days, while in other sites the sampling extended over several weeks, with each individual sampling day covering only a few leaves or species. In those cases, an adequate representation of the multi-species canopy is achieved only when data from all sampling days are considered. The example shown in Figure 7 corresponds to a site whose leaf sampling campaign lasted almost one month. During this time, the carotenoid reflectance index (CRI_550_) exhibited pronounced fluctuations throughout the sampling period (Figure 7a). CRI_550_ values ranged from 6.5 to 14, representing a variation of nearly 50% the overall range found across all sites (see full cross-site CRI_550_ range in Figure 4b and 4e). Interestingly, the satellite images of the same site for those dates (Figure 7b-c) showed no visible differences in canopy structure, disturbance, or atmospheric conditions, and such within-season VI fluctuations may also contribute to the mismatch between satellite data and our field leaf trait estimates.

**Figure 7.**
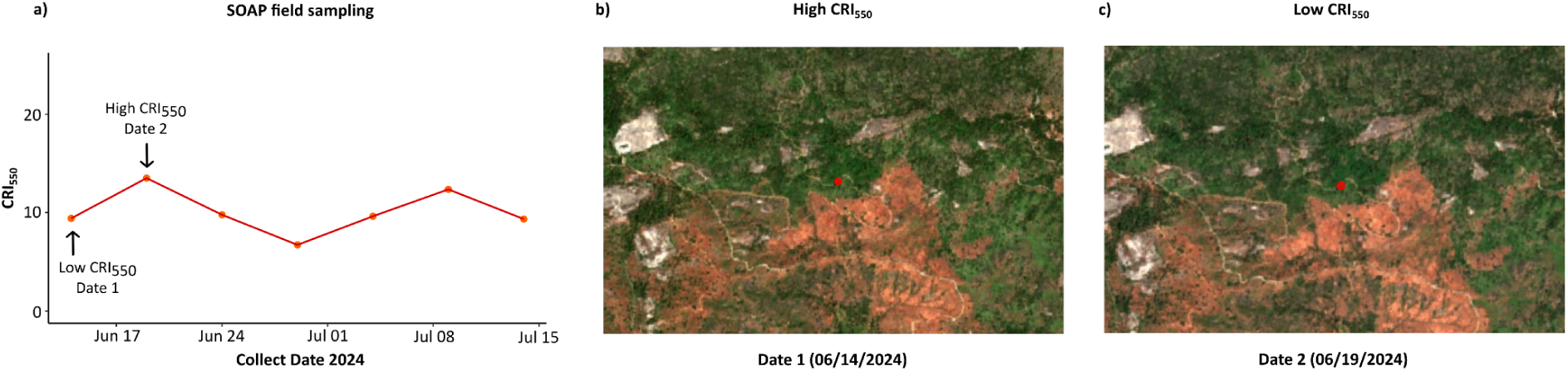
Example of wide temporal variation in CRI_550_ that occurred during a prolonged leaf traits field sampling. In this NEON site, carotenoid reflectance index (CRI₅₅₀) values fluctuated strongly across different sampling dates within the same campaign (a), despite no detectable changes in canopy (e.g., LAI) or atmospheric conditions (e.g., no clouds were present) in satellite images from those dates (b, c).

## 4. DISCUSION

Our model-based sensitivity analysis revealed that the evaluated VIs were able to detect variation in their target traits—leaf water content, carotenoids, or the chlorophyll/carotenoid ratio—only to a limited extent. In contrast, LAI consistently emerged as the most influential factor across all VIs, often surpassing the contribution of the leaf traits these VIs were intended to estimate. When analyzed with ground truth and satellite data, however, these indices were able to capture only a minimal fraction of the observed variation in foliar traits or even none in some cases. This discrepancy between model- and observation-based findings indicates that in real conditions, the signal of water and carotenoid indices, besides being is largely driven by LAI and other structural traits such as LMA, is further influenced by factors not often included in the model-based analyses, such as atmospheric conditions and even issues related to field sampling protocols.

Our findings confirm that LAI is the main factor influencing VIs variability, even surpassing the biochemical traits they are meant to represent. This finding is consistent with previous model-based studies, which show a large influence of LAI and other structural traits on general-purpose VIs (e.g., NDVI, EVI) and indices for water and chlorophyll (Morcillo-Pallarés et al. 2019; Verrelst et al. 2015; Hinojo-Hinojo & Goulden 2021; Tian et al. 2025, Hinojo-Hinojo et al. 2024). It is well known that VIs tend to saturate at moderate to high LAI values and to be strongly influenced by soil reflectance at low LAI values (Tian et al 2025). This soil influence may explain the reduced capacity of VIs to capture leaf water and carotenoid content at low LAI values, shown in our model-based analyses (Figure 2a, d and g). More importantly, our model-based analyses detected an interesting opportunity: when VIs saturate to LAI, most VIs variation is driven by leaf water and carotenoid content (Figure 2b-c, e-f, h-i). This implies that, if atmospheric noise, other unwanted signals and potential sampling biases could be ruled out (see below), VIs could be a great tool to monitor leaf water and carotenoid contents in forest and other high LAI ecosystems. Lastly, the large influence of LAI over carotenoid and chlorophyll/carotenoid ratio indices raises questions over the mechanisms behind the strong relationship between these indices and vegetation primary production and phenology across ecosystems (e.g. Wang et al 2023): it is possible that such relationships are not solely driven by stress responses associated with carotenoids (Gamon et al 2016), but also due to those VIs strong relationship to LAI (Figures 2d and g, and 4f).

Although RTMs are widely used and are valuable tools for studying VIs sensitivity factors, recent work has highlighted that they still present important limitations that could help explain the patterns of limited VIs sensitivity and strong dominance of LAI we observed in our results. Wang et al. (2025) noted that many RTMs remain overly simplified, which limits their ability to reproduce the full spectral variability of plant leaf and canopy optical properties. For example, the influence of litter and stems over surface reflectance is rarely included in the most widely-used RTM (Jacquemoud et al 2009, Ligot et al 2014), but can drive substantial variation in surface reflectance in certain ecosystem types (Asner 1998). This simplification suggests that, while RTM-based sensitivity analyses are useful for identifying general trends, their predictive accuracy under diverse ecosystems, species composition and vegetation growing dynamics may be more limited than expected. These limitations of RTMs may have contributed to the mismatch in our findings of VIs sensitivity in our model-based vs field-based assessments.

Another factor that introduces uncertainty in our findings comes from atmospheric conditions. Coluzzi et al. (2018) noted that the presence of thin clouds or cirrus can introduce significant noise into reflectance signals by altering radiation transmission and reducing the accuracy of surface parameter detection. There exist mature algorithms for performing quality control for atmospheric conditions in the multispectral sensors used in this study.

However, some clouds are so thin and/or opaque that they can remain undetected by such algorithms. In our study, we documented a high sensitivity of CRI_550_ to this type of cloud, much higher than NDVI (Figure 6). This could indicate that part of the CRI_550_ captured variability is not due to physiological changes but rather to atmospheric conditions that are difficult to filter out with the products used to obtain satellite images.

Regarding ground truth observations, data quality can also vary due to sampling biases. At several NEON sites, understory vegetation was neither consistently sampled nor its presence consistently noted in the data. In some of the sites, understory presence is quite significant and therefore potentially contributes significantly to the spectral signature of the plot (Hinojo-Hinojo et al. 2024). Previous studies have noted that the presence of understory can markedly alter canopy reflectance, especially in open forests where light penetrates into deeper layers (Eriksson et al. 2006). This could explain why in our data we observed great data variability in sites such as RMNP, YELL, SOAP, or LENO, where plots dominated by tree canopies coexist with others located in more open areas covered mainly by herbaceous vegetation (Figure 5). In sites where understory vegetation was abundant but not well sampled, the vegetation trait data may not be properly representing the functional and optical properties of the canopy, weakening the association between the VIs and the ground truth data. This sampling bias of forest understory is still common (Landuyt et al 2019), and adequate protocols for characterizing it should be included in ecological monitoring efforts, and appropriate ways to weight their contribution need to be developed, especially in those tailored for ground-truthing remote sensing approaches.

Another limitation we found in the field data is related to the duration of the sampling campaigns. At several NEON sites, leaf trait sampling extended beyond a few days, lasting for several weeks and in some cases close to a month. This implies that the measurements may be unintentionally incorporating more variability due to physiological responses to the environment and the phenological stage of sampled vegetation. Recent studies have shown that LWC can vary over very short time windows, even within days (Junttila et al. 2022).

Similarly, carotenoids undergo rapid changes during phenological transitions, with detectable shifts occurring within days to weeks (Wong et al. 2022). In our results, we detected pronounced temporal fluctuations CRI_550_ in sites with prolonged sampling campaigns and those fluctuations occurred without noticeable changes in canopy structure or atmospheric conditions (Figure 7). These fluctuations may reflect natural carotenoid dynamics (and potentially other functional traits) occurring within the sampling period. Consequently, prolonged field campaigns can introduce temporal variation, as measurements collected over several weeks may represent different vegetation phenological stages, thereby weakening the association between field and satellite observations.

### 4.1. Future directions

Based on the limitations identified in our study, we suggest some adjustments that could improve the reliability and generalization of VIs in plant studies. In the case of RTMs, it may be necessary to seek radiative transfer schemes that better integrate the natural dynamics of vegetation and seasonal changes in foliar traits, other potentially important factors such as litter and stems contribution to surface reflectance, and even noise from atmospheric conditions. It may also be useful to integrate RTMs into simulation frameworks that explicitly replicate common sampling strategies and biases, allowing for the identification of VIs or metrics less influenced by these real-world limitations. For satellite sensor data, improved atmospheric correction products would help minimize the noise caused by thin clouds and other atmospheric interferences. Regarding ground truth data collection in open forests, it is important not only to consider tree species but also to include understory vegetation and non-tree species, ensuring a more complete representation of the canopy spectral signal.

Furthermore, it is advisable to standardize the timing of sampling campaigns based on phenological stages, conducting shorter surveys, if possible, more than once a year to reduce phenological inconsistencies. It may also be useful to keep developing other innovative resources, such as hyperspectral data and methods that are best suited to capture the leaf traits signal with greater precision, and machine learning models, to further inform the improvement and design of VIs, including multispectral (Lamour et al. 2025). Although these recommendations will not completely eliminate the current limitations, they can contribute to improving the reliability of multispectral proxies, thereby opening new possibilities for monitoring vegetation at large spatial and temporal scales.

Finally, our findings indicate that multispectral VIs have limited ability to capture foliar water and carotenoid contents across different ecosystems, as their signals are mainly dominated by LAI, and their performance is further reduced by atmospheric conditions and even field sampling strategies. Addressing these limitations through strategies that improve field sampling and the integration of advanced methodologies across spatial scales (e.g., leaf-canopy-ecosystem) to complement these analyses will be key to strengthening future vegetation monitoring studies under an increasingly stressful climate.

## 5. CONCLUSIONS

Our analyses revealed that under simulated conditions, multispectral VIs were able to capture only a small fraction of the variation in leaf water and carotenoids content, and the chlorophyll/carotenoid ratio. The indices that performed best were NDWI for water content, CRI_550_ for carotenoids, and CCI for the chlorophyll/carotenoid ratio, particularly in high LAI canopies. All indices analyzed showed a predominant sensitivity to LAI, but sensitivity to LAI decreases and sensitivity to leaf water content increases at LAI > 2. These findings reveal that if atmospheric noise, other unwanted signals and potential sampling biases could be ruled out, multispectral VIs could be a great tool to monitor leaf water and carotenoid contents in forest and other high LAI ecosystems

When evaluating the performance of the VIs against ground truth and satellite data, the indices showed almost no sensitivity to their target biochemical traits (≤6% of the variation explained). In addition, the influence of LAI was even stronger, as LAI greatly dominated the VIs sensitivity.

On the other hand, we found factors that can be the main causes of why the VIs were not associated with the pigments they are meant to predict:

● Data collection and image artifacts, in particular, the limitations of RTMs in accounting for the diversity of vegetation dynamics and conditions
● Nearly imperceptible atmospheric noise such as thin clouds and cirrus that remain uncorrected even after applying filters to satellite images
● Bias in field sampling, such as the lack of representation of understory and non-tree species
● Prolonged field campaigns that introduce temporal variability due to phenology Strategies we recommend to improve the applicability of VIs:
● Avoid using multispectral VIs to estimate foliar water and pigment content in ecosystems where canopy vegetation is sparse (e.g. LAI < 2)
● Prioritize using images with minimal cloud presence and use advanced atmospheric quality control products
● Conduct a representative sampling of plant species, including understory and non-tree vegetation
● Standardize the duration of field sampling campaigns
● Integrate complementary tools and approaches to strengthen and support analyses based on VIs

Exploring the limitations of the analyses carried out helped us identify and propose strategies to improve vegetation stress monitoring through leaf water and carotenoid contents and chlorophyll/carotenoid ratio. It is necessary to refine the techniques with which we analyze vegetation status in order to obtain more precise information at large spatial and temporal scales, which can help us develop effective strategies for the conservation of vegetation and the assessment of climate change effects on plant function or the productivity of crops.

## DATA AND CODE STATEMENT

The NEON data used are publicly available at the NEON data portal located at https://www.neonscience.org/data. For the plant foliar trait product, we used both release and provisional data, and we only used release data for the vegetation structure product, both products downloaded on February 14 of 2025. The specific product versions used are available at https://doi.org/10.17605/OSF.IO/4XCJR. The codes used for processing and analysing the data from this manuscript are available at: https://gitlab.com/hinojohinojo/water_carotenoids_indices.git

## ACKNOWLEDGMENTS

This work was supported by the U.S. National Science Foundation’s Biodiversity on a Changing Planet (BoCP) Program [grant numbers: BoCP-2416164, BoCP-2225076], and the Secretaría de Ciencia, Humanidades, Tecnología e Inovación (SECIHTI) thought the program Apoyos a Laboratorios Nacionales [grant number: ApoyoLN-2025-C-1]. CHH received a repatriation fellowship from Consejo Nacional de Ciencia y Tecnología (CONACyT)’s 2022 repatriacion program [agreement number: I1200/320/2022 MOD.ORD./09/2022]. GMM thanks Secretaría de Ciencia, Humanidades, Tecnología e Inovación (SECIHTI) for a national postdoctoral fellowship. CARZ acknowledges support from the LOEWE Professorship for Plant Breeding at Hochschule Geisenheim University funded by the Hessian Ministry of Higher Education, Research and the Arts in Hesse, Germany while working on this manuscript. The NEON datasets used come from work supported by the National Ecological Observatory Network (NEON), a program sponsored by the U.S. National Science Foundation (NSF) and operated under cooperative agreement by Battelle. We acknowledge Grecia Fernanda Rivas Ríos for her support on the processing of leaf traits and statistical plot trait distributions, and León Arturo Jara Sarracino and Ramón Ángel López Castro for their involvement in gathering and processing leaf area index data for the NEON field sites.

## AUTHOR CONTRIBUTIONS

**MGMB:** Conceptualization, Data curation, Formal analysis, Investigation, Methodology, Software, Visualization, Writing – original draft, Writing - Review and Editing

**CARZ:** Conceptualization, Methodology, Investigation, Supervision, Writing - Review and Editing

**CTO:** Supervision, Writing - Review and Editing

**BJE:** Funding acquisition, Supervision, Writing - Review and Editing

**AF:** Funding acquisition, Supervision, Writing - Review and Editing

**BM:** Writing - Review and Editing

**GMM:** Writing - Review and Editing

**JRRL:** Resources, Supervision

**LS:** Writing - Review and Editing

**CHH:** Conceptualization, Data curation, Formal analysis, Funding acquisition, Investigation, Methodology, Project administration, Software, Supervision, Visualization, Writing – original draft, Writing - Review and Editing

## REFERENCES

1. Asner, G.P., 1998. Biophysical and biochemical sources of variability in canopy reflectance. Remote Sens. Environ. 64, 234–253. 10.1016/S0034-4257(98)00014-5

2. Boving, I., Allen, J., Brodrick, P., Chadwick, K., Trugman, A., Anderegg, L., 2025. The unstable relationship between drought status and leaf water content complicates the remote sensing of tree drought stress. Glob. Change Biol. 31, e70188. 10.1111/gcb.70188

3. Coluzzi, R., Imbrenda, V., Lanfredi, M., Simoniello, T., 2018. A first assessment of the Sentinel-2 Level 1-C cloud mask product to support informed surface analyses. Remote Sens. Environ. 217, 426–443. 10.1016/j.rse.2018.08.009

4. Dall’Osto, L., Bassi, R., Ruban, A., 2014. Photoprotective mechanisms: Carotenoids. Plastid Biol. 5, 393–435. 10.1007/978-1-4939-1136-3_15

5. Eriksson, H.M., Eklundh, L., Kuusk, A., Nilson, T., 2006. Impact of understory vegetation on forest canopy reflectance and remotely sensed LAI estimates. Remote Sens. Environ. 103, 408–418. 10.1016/j.rse.2006.04.005

6. Fahad, S., Bajwa, A.A., Nazir, U., Anjum, S.A., Farooq, A., Zohaib, A., Sadia, S., Nasim, W., Adkins, S., Saud, S., Ihsan, M.Z., Alharby, H., Wu, C., Wang, D., Huang, J., 2017. Crop production under drought and heat stress: Plant responses and management options. Front. Plant Sci. 8, 1147. 10.3389/fpls.2017.01147.

7. Féret, J.-B., François, C., Asner, G.P., Gitelson, A.A., Martin, R.E., Bidel, L.P.R., Ustin, S.L., le Maire, G., Jacquemoud, S., 2008. PROSPECT-4 and 5: Advances in the leaf optical properties model separating photosynthetic pigments. Remote Sens. Environ. 112, 3030–3043. 10.1016/j.rse.2008.02.012

8. Gamon, J., Huemmrich, K., Wong, C., Ensminger, I., Garrity, S., Hollinger, D., Noormets, A., Peñuelas, J., 2016. A remotely sensed pigment index reveals photosynthetic phenology in evergreen conifers. Proc. Natl. Acad. Sci. U.S.A. 113, 13087–13092. 10.1073/pnas.1606162113

9. Gao, B.-C., 1996. NDWI—a normalized difference water index for remote sensing of vegetation liquid water from space. Remote Sens. Environ. 58, 257–266. 10.1016/S0034-4257(96)00067-3.

10. Hais, M., Hellebrandová, K.N., Šrámek, V., 2019. Potential of Landsat spectral indices in regard to the detection of forest health changes due to drought effects. *J*. For. Sci. 65, 70–78. 10.17221/137/2018-JFS

11. Hammond, W., Williams, A., Abatzoglou, J., Adams, H., Klein, T., López, R., Sáenz-Romero, C., Hartmann, H., Breshears, D., & Allen, C. (2022). Global field observations of tree die-off reveal hotter-drought fingerprint for Earth’s forests. Nature Communications, 13(1), 1761. 10.1038/s41467-022-29289-2

12. Hernández-Clemente, R., Navarro-Cerrillo, R.M., Zarco-Tejada, P.J., 2012. Carotenoid content estimation in a heterogeneous conifer forest using narrow-band indices and PROSPECT+DART simulations. Remote Sens. Environ. 127, 298–315. 10.1016/j.rse.2012.09.014

13. Hinojo-Hinojo, C., Goulden, M.L., 2020a. Plant traits help explain the tight relationship between vegetation indices and gross primary production. Remote Sens. 12(9). 10.3390/RS12091405

14. Hinojo-Hinojo, C., Goulden, M.L., 2020b. A compilation of canopy leaf inclination angle measurements across plant species and biome types [dataset]. Dryad, v1. 10.7280/D1T97H

15. Hinojo-Hinojo, C., Bohner, T., Chacón-Labella, J., Chmurzynski, A., Falco, N., Goulden, M.L., Hemingway, B.L., Merow, C., Nikolopoulos, E.I., Wang, J.A., Wainwright, H., Frazier, A.E., Enquist, B.J., n.d. Global shift in a key plant trait indicates a change in biosphere function. OSF Preprints. 10.31219/osf.io/zs4fp

16. Homolová, L., Malenovský, Z., Clevers, J.G.P.W., García-Santos, G., Schaepman, M.E., 2013. Review of optical-based remote sensing for plant trait mapping. Ecol. Complex. 15, 1–16. 10.1016/J.ECOCOM.2013.06.003

17. Huete, A., Justice, C., Liu, H., 1994. Development of vegetation and soil indices for MODIS-EOS. Remote Sens. Environ. 49, 224–234. 10.1016/0034-4257(94)90018-3

18. Jacquemoud, S., Verhoef, W., Baret, F., Bacour, C., Zarco-Tejada, P., Asner, G., François, C., Ustin, S., 2009. PROSPECT + SAIL models: A review of use for vegetation characterization. Remote Sens. Environ. 113(Suppl. 1), S56–S66. 10.1016/j.rse.2008.01.026

19. Ji, L., Zhang, L., Wylie, B.K., Rover, J., 2011. On the terminology of the spectral vegetation index (NIR − SWIR)/(NIR + SWIR). Int. J. Remote Sens. 32, 6901–6909. 10.1080/01431161.2010.510811

20. Jones, H.G., 2014. Plants and microclimate: A quantitative approach to environmental plant physiology, third ed. Cambridge University Press, Cambridge. 10.1017/CBO9780511845727

21. Junttila, S., Junttila, S., Hölttä, T., Saarinen, N., Kankare, V., Yrttimaa, T., Hyyppä, J., Vastaranta, M., 2022. Close-range hyperspectral spectroscopy reveals leaf water content dynamics. Remote Sens. Environ. 277, 113071. 10.1016/j.rse.2022.113071

22. Landuyt, D., de Lombaerde, E., Perring, M., Hertzog, L., Ampoorter, E., Maes, S., de Frenne, P., Ma, S., Proesmans, W., Blondeel, H., Sercu, B., Wang, B., Wasof, S., Verheyen, K., 2019. The functional role of temperate forest understorey vegetation in a changing world. Glob. Change Biol. 25, 3625–3641. 10.1111/gcb.14756

23. Lamour, J., Serbin, S., Rogers, A., Acebron, K., Ainsworth, E., Albert, P., Alonzo, M., Anderson, J., Atkin, O., Barbier, N., Barnes, M., Bernacchi, C., Besson, N., Burnett, A., Caplan, J., Chave, J., Cheesman, A., Clocher, I., Coast, O., Zhao, Y., 2025. The Global Spectra-Trait Initiative: A database of paired leaf spectroscopy and functional traits associated with leaf photosynthetic capacity. Earth Syst. Sci. Data Discuss. 2025, 1–34. 10.5194/essd-2025-213

24. Lehnert, L.W., Meyer, H., Obermeier, W.A., Silva, B., Regeling, B., Bendix, J., 2019. Hyperspectral Data Analysis in R: The hsdar Package. J. Stat. Softw. 89(12), 1–23. 10.18637/jss.v089.i12

25. Li, X., Zhu, B., Li, S., Liu, L., Song, K., Liu, J., 2025. A comprehensive review of crop chlorophyll mapping using remote sensing approaches: achievements, limitations, and future perspectives. Sensors 25, 2345. 10.3390/s25082345

26. Li, S., Fang, H., 2025. Mapping global leaf inclination angle (LIA) based on field measurement data. *Earth Syst*. Sci. Data 17, 1347–1366. 10.5194/essd-17-1347-2025

27. Ligot, G., Balandier, P., Courbaud, B., Claessens, H., 2014. Forest radiative transfer models: Which approach for which application? Can. J. For. Res. 44, 391–403. 10.1139/cjfr-2013-0494

28. Maitner, B., Halbritter, A., Telford, R., Strydom, T., Chacon, J., Lamanna, C., Sloat, L., Kerkhoff, A., Messier, J., Rasmussen, N., Pomati, F., Merz, E., Vandvik, V., Enquist, B., 2023. Bootstrapping outperforms community-weighted approaches for estimating the shapes of phenotypic distributions. Methods Ecol. Evol. 14, 2592–2610. 10.1111/2041-210X.14160

29. Morcillo-Pallarés, P., Rivera-Caicedo, J., Belda, S., de Grave, C., Burriel, H., Moreno, J., Verrelst, J., 2019. Quantifying the robustness of vegetation indices through global sensitivity analysis of homogeneous and forest leaf-canopy radiative transfer models. Remote Sens. 11, 2418. 10.3390/rs11202418

30. Murchie, E., Lawson, T., 2013. Chlorophyll fluorescence analysis: A guide to good practice and understanding some new applications. J. Exp. Bot. 64, 3983–3998. 10.1093/jxb/ert208

31. Oksanen, J., Simpson, G., Blanchet, F., Kindt, R., Legendre, P., Minchin, P., O’Hara, R., Solymos, P., Stevens, M., Szoecs, E., Wagner, H., Barbour, M., Bedward, M., Bolker, B., Borcard, D., Carvalho, G., Chirico, M., De Cáceres, M., Durand, S., Evangelista, H., FitzJohn, R., Friendly, M., Furneaux, B., Hannigan, G., Hill, M., Lahti, L., McGlinn, D., Ouellette, M., Ribeiro Cunha, E., Smith, T., Stier, A., ter Braak, C., Weedon, J., Borman, T., 2025. vegan: Community ecology package (R package version 2.6-10) [software]. https://CRAN.R-project.org/package=vegan.

32. Ollinger, S.V., 2011. Sources of variability in canopy reflectance and the convergent properties of plants. New Phytol. 189, 375–394. 10.1111/j.1469-8137.2010.03536.x

33. Pérez-Harguindeguy, N., Díaz, S., Garnier, E., Lavorel, S., Poorter, H., Jaureguiberry, P., Bret-Harte, M.S., Cornwell, W.K., Craine, J.M., Gurvich, D.E., Urcelay, C., Veneklaas, E.J., Reich, P.B., Poorter, L., Wright, I.J., Ray, P., Enrico, L., Pausas, J.G., de Vos, A.C., Cornelissen, J.H.C., 2013. New handbook for standardised measurement of plant functional traits worldwide. Aust. J. Bot. 61, 167–234. 10.1071/BT12225

34. Saltelli, A., Annoni, P., Azzini, I., Campolongo, F., Ratto, M., Tarantola, S., 2010. Variance based sensitivity analysis of model output. Design and estimator for the total sensitivity index. Comput. Phys. Commun. 181, 259–270. 10.1016/j.cpc.2009.09.018

35. Schimel, D., Schneider, F.D., 2019. Flux towers in the sky: Global ecology from space. New Phytol. 224, 570–584. 10.1111/nph.15934

36. Simkin, A., Kapoor, L., Doss, C., Hofmann, T., Lawson, T., Ramamoorthy, S., 2022. The role of photosynthesis related pigments in light harvesting, photoprotection and enhancement of photosynthetic yield in plants. Photosynth. Res. 152, 23–42. 10.1007/s11120-021-00892-6

37. Sims, D., Gamon, J., 2002. Relationships between leaf pigment content and spectral reflectance across a wide range of species, leaf structures and developmental stages. Remote Sens. Environ. 81, 337–354. 10.1016/S0034-4257(02)00010-X

38. Sun, T., Rao, S., Zhou, X., Li, L., 2022. Plant carotenoids: Recent advances and future perspectives. Mol. Hortic. 2, 23. 10.1186/s43897-022-00023-2

39. Tian, X., Jia, X., Da, Y., Liu, J., Ge, W., 2025. Evaluating the sensitivity of vegetation indices to leaf area index variability at individual tree level using multispectral drone acquisitions. Agric. For. Meteorol. 364, 110896. 10.1016/j.agrformet.2025.110441

40. Verhoef, W., 1984. Light scattering by leaf layers with application to canopy reflectance modeling: The SAIL model. Remote Sens. Environ. 16, 125–141. 10.1016/0034-4257(84)90057-9

41. Verrelst, J., Rivera, J., Moreno, J., 2015. ARTMO’s Global Sensitivity Analysis (GSA) toolbox to quantify driving variables of leaf and canopy radiative transfer models. EARSeL eProc. Spec. Issue 2, 1–11. 10.12760/02-2015-2-01

42. Vidican, R., Mălinaș, A., Ranta, O., Moldovan, C., Marian, O., Ghețe, A., Ghișe, C.R., Popovici, F., Cătunescu, G.M., 2023. Using remote sensing vegetation indices for the discrimination and monitoring of agricultural crops: A critical review. Agronomy 13, 12. 10.3390/agronomy13123040

43. Wang, R., Bowling, D.R., Gamon, J.A., Smith, K.R., Yu, R., Hmimina, G., Ueyama, M., Noormets, A., Kolb, T.E., Richardson, A.D., Bourque, C.P.A., Bracho, R., Blanken, P.D., Black, T.A., Arain, M.A., 2023. Snow-corrected vegetation indices for improved gross primary productivity assessment in North American evergreen forests. Agric. For. Meteorol. 340, 109600. 10.1016/J.AGRFORMET.2023.109600

44. Wang, Y., Braghiere, R.K., Fischer, W.W., Yao, Y., Shen, Z., Schneider, T., Bloom, A.A., Schimel, D., Croft, H., Winkler, A.J., Reichstein, M., Frankenberg, C., 2025. Impacts of leaf traits on vegetation optical properties in Earth system modeling. Nat. Commun. 16, 4968. 10.1038/s41467-025-60149-x

45. Weintraub-Leff, S., 2024. NEON user guide to plant foliar traits (DP1.10026.001), Revision G. National Ecological Observatory Network, Boulder, CO.

46. Xiao, Y., Zhao, W., Zhou, D., Gong, H., 2014. Sensitivity analysis of vegetation reflectance to biochemical and biophysical variables at leaf, canopy, and regional scales. IEEE Trans. Geosci. Remote Sens. 52, 4014–4024. 10.1109/TGRS.2013.2278838

47. Xu, H., Sun, H., Xu, Z., Wang, Y., Zhang, T., Wu, D., Gao, J.H., 2025. kNDMI: A kernel normalized difference moisture index for remote sensing of soil and vegetation moisture. Remote Sens. Environ. 319, 114621. 10.1016/J.RSE.2025.114621

48. Xue, J., Su, B., 2017. Significant remote sensing vegetation indices: A review of developments and applications. J. Sens. 2017, 1–17. 10.1155/2017/1353691

49. Yan, K., Gao, S., Yan, G., Ma, X., Chen, X., Zhu, P., Li, J., Gao, S., Gastellu-Etchegorry, J.P., Myneni, R.B., Wang, Q., 2025. A global systematic review of the remote sensing vegetation indices. Int. J. Appl. Earth Obs. Geoinf. 139, 104951. 10.1016/j.jag.2025.104560

50. Yang, X., Lu, M., Wang, Y., Wang, Y., Liu, Z., Chen, S., 2021. Response mechanism of plants to drought stress. Horticulturae 7, 85. 10.3390/horticulturae7030050

51. Yasir, Q., Zhang, Z., Ren, J., Wang, G., Naveed, M., Jahangir, Z., Rahman, A., 2024. Spectral index for estimating leaf water content across diverse plant species using multiple viewing angles. J. Appl. Remote Sens. 18, 048501. 10.1117/1.jrs.18.042603

52. Zandalinas, S., Mittler, R., 2022. Plant responses to multifactorial stress combination. New Phytol. 234, 1161–1167. 10.1111/nph.18087

