## Supplementary material for "Challenges and opportunities in detecting leaf water and carotenoid content across biomes from satellite multispectral indices"

**This file includes:**

Table S1 and Figures S1 to S9.

**Table S1.** NEON sites used in this study. The table includes the site name, official abbreviation, ecosystem classification, and the year of biochemical/physiological trait data collection. Satellite information was extracted using the geographic coordinates of each site

| ID | Site | Abbreviation | Type | Collect Year |
|---|---|---|---|---|
| 1 | Abby Road | ABBY | Evergreen forest | 2022 |
| 2 | Utqiaġvik | BARR | Herbaceous | 2022 |
| 3 | Bartlett Experimental Forest | BART | Deciduous broadleaf forest | 2022 |
| 4 | Caribou-Poker Creeks Research Watershed | BONA | Deciduous forest | 2024 |
| 5 | Lyndon B. Johnson National Grassland | CLBJ | Deciduous broadleaf forest | 2024 |
| 6 | Central Plains Experimental Range | CPER | Grassland/herbaceous | 2021 |
| 7 | Dakota Coteau Field | DCFS | Grassland/herbaceous | 2019 |
| 8 | Delta Junction | DEJU | Evergreen conifer forest | 2023 |
| 9 | Dead Lake | DELA | Deciduous broadleaf forest | 2024 |
| 10 | Great Smoky Mountains National Park | GRSM | Deciduous broadleaf forest | 2021 |
| 11 | Guánica Forest | GUAN | Evergreen broadleaf forest | 2022 |
| 12 | Harvard Forest & Quabbin Watershed | HARV | Mixed forest | 2024 |
| 13 | The Jones Center At Ichauway | JERC | Evergreen conifer forest | 2024 |
| 14 | Jornada Experimental Range | JORN | Herbaceous/grassland | 2022 |
| 15 | Konza Prairie Agroecosystem | KONA | Grassland/herbaceous | 2019 |
| 16 | Konza Prairie Biological Station | KONZ | Grassland/herbaceous | 2022 |
| 17 | Lajas Experimental Station | LAJA | Pasture/Hay | 2020 |
| 18 | Lenoir Landing | LENO | Deciduous broadleaf forest | 2023 |
| 19 | Mountain Lake Biological Station | MLBS | Deciduous broadleaf forest | 2023 |
| 20 | Northern Great Plains Research Laboratory | NOGP | Grassland/herbaceous | 2020 |
| 21 | Marvin Klemme Range Research Station | OAES | Shrub/Scrub | 2023 |

| | | | | |
|---|---|---|---|---|
| 22 | Onaqui | ONAQ | Shrub/Scrub | 2021 |
| 23 | Oak Ridge | ORNL | Deciduous broadleaf forest | 2022 |
| 24 | Ordway-Swisher Biological Station | OSBS | Evergreen forest | 2021 |
| 25 | Pu'u Maka'ala Natural Area Reserve | PUUM | Evergreen forest | 2019 |
| 26 | Rocky Mountains | RMNP | Evergreen forest | 2020 |
| 27 | Smithsonian Conservation Biology Institute | SCBI | Deciduous broadleaf forest | 2022 |
| 28 | Smithsonian Environmental Research Center | SERC | Deciduous broadleaf forest | 2021 |
| 29 | Soaproot Saddle | SOAP | Evergreen conifer forest | 2024 |
| 30 | Steigerwaldt-Chequamegon | STEI | Deciduous forest | 2022 |
| 31 | North Sterling | STER | Cultivated crops | 2022 |
| 32 | Talladega National Forest | TALL | Evergreen forest | 2021 |
| 33 | Lower Teakettle | TEAK | Evergreen forest | 2021 |
| 34 | Toolik Field Station | TOOL | Herbaceous | 2022 |
| 35 | Treehaven | TREE | Mixed forest | 2020 |
| 36 | KU Field Station | UKFS | Deciduous forest | 2023 |
| 37 | University of Notre Dame Environmental Research Center | UNDE | Deciduous forest | 2024 |
| 38 | Chase Lake National Wildlife Refuge | WOOD | Grassland/herbaceous | 2021 |
| 39 | Yellowstone National Park | YELL | Evergreen forest | 2023 |

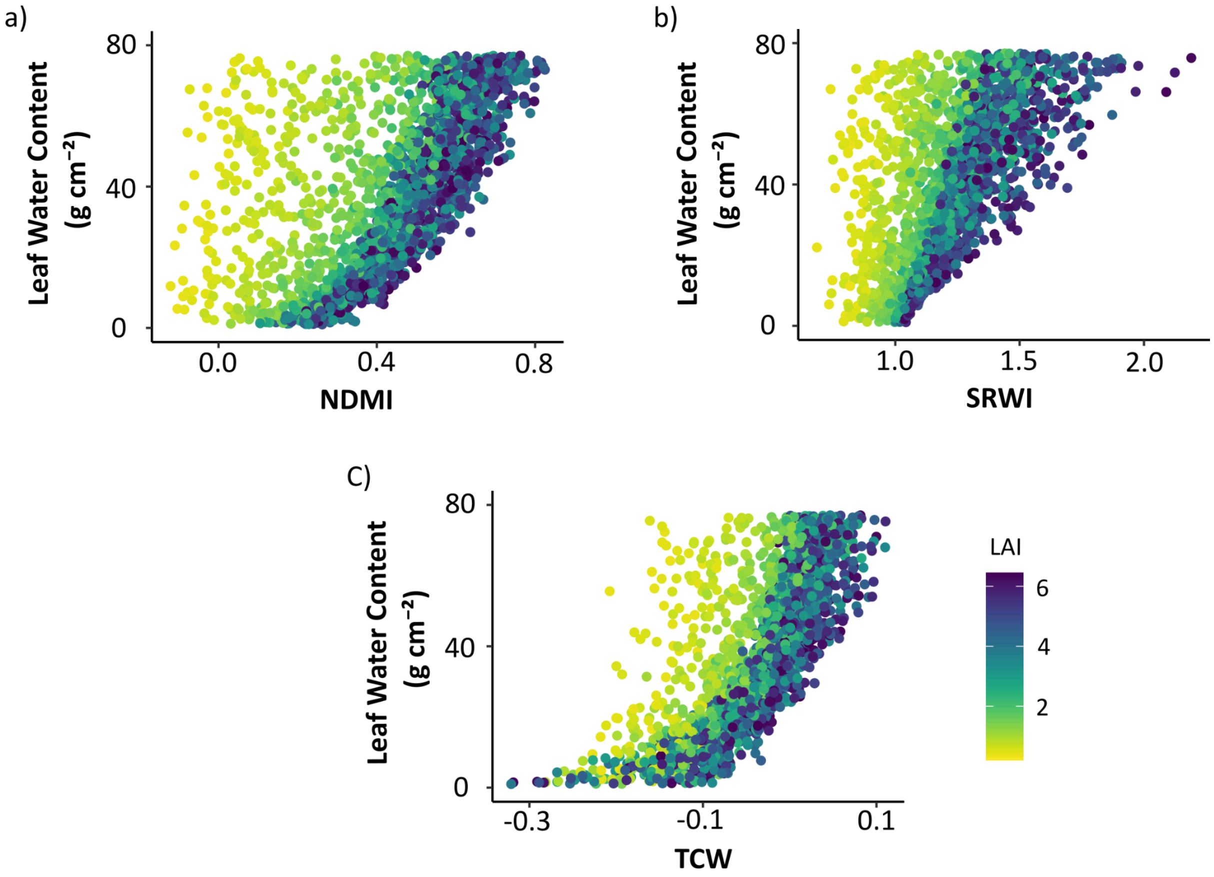


**Figure S1.** Relationships between three multispectral indices associated with water content — NDMI, SRWI y TCW — and leaf water content simulated with the PROSAIL model. Each point represents a unique combination of parameter values generated with the PROSAIL model, i.e., a canopy with particular values of leaf and canopy traits, as well as environmental and sun/sensor geometry conditions. The color gradient (yellow to dark blue) indicates increasing leaf area index (LAI) values, from sparse to dense canopies. All indices are shown across the full LAI range (0.2–7). The strong dependence of index variation on LAI highlights the influence of canopy structure on the spectral signal associated with water.

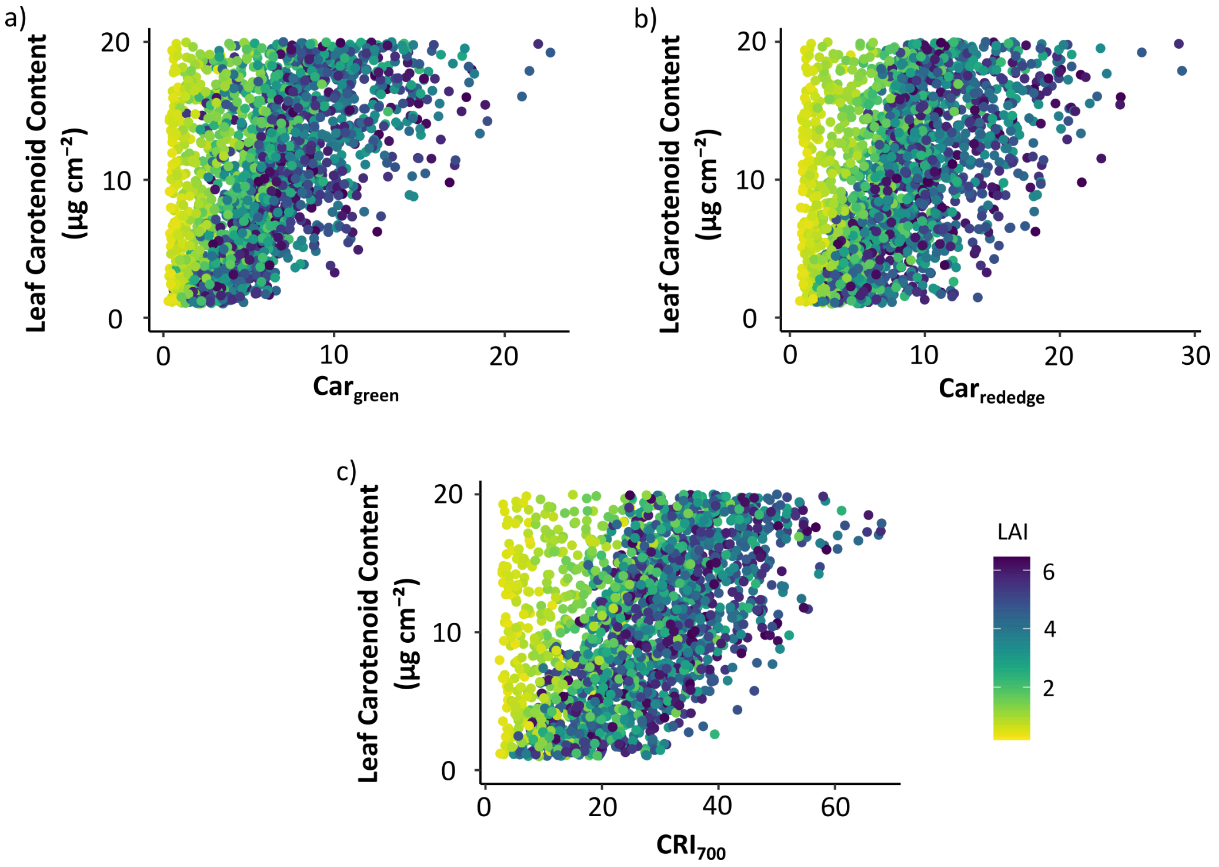


**Figure S2.** Relationships between three multispectral indices associated with carotenoid content — $Car_{green}$, $Car_{rededge}$, and $CRI_{700}$ — and leaf carotenoid content simulated with the PROSAIL model. Each point represents a unique combination of parameter values generated with the PROSAIL model, i.e., a canopy with particular values of leaf and canopy traits, as well as environmental and sun/sensor geometry conditions. The color gradient (yellow to dark blue) indicates increasing leaf area index (LAI) values, from sparse to dense canopies. All indices are shown across the full LAI range (0.2–7). The strong dependence of index variation on LAI highlights the influence of canopy structure on the spectral signal associated with carotenoids.

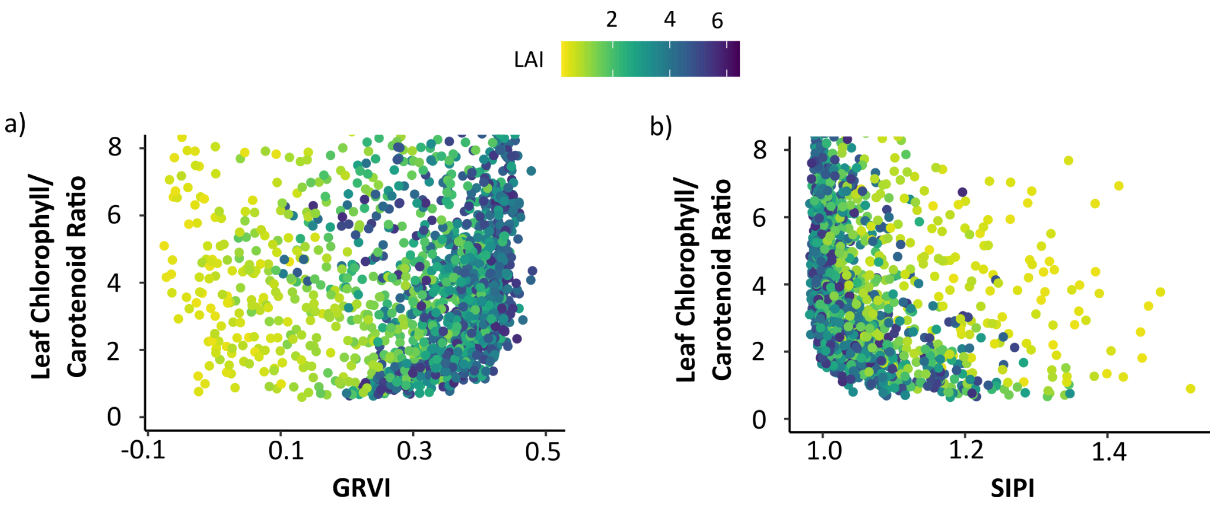


**Figure S3.** Relationships between two multispectral indices associated with the chlorophyll/carotenoid ratio -SIPI and GRVI- and the leaf chlorophyll/carotenoid ratio simulated with the PROSAIL model. Each point represents a unique combination of parameter values generated with the PROSAIL model, i.e., a canopy with particular values of leaf and canopy traits, as well as environmental and sun/sensor geometry conditions. The color gradient (yellow to dark blue) indicates increasing leaf area index (LAI) values, from sparse to dense canopies. All indices are shown across the full LAI range (0.2–7). The weak relationships observed and their dependence on LAI reflect the stronger influence of canopy structure on the spectral signal compared with the chlorophyll/carotenoid ratio.

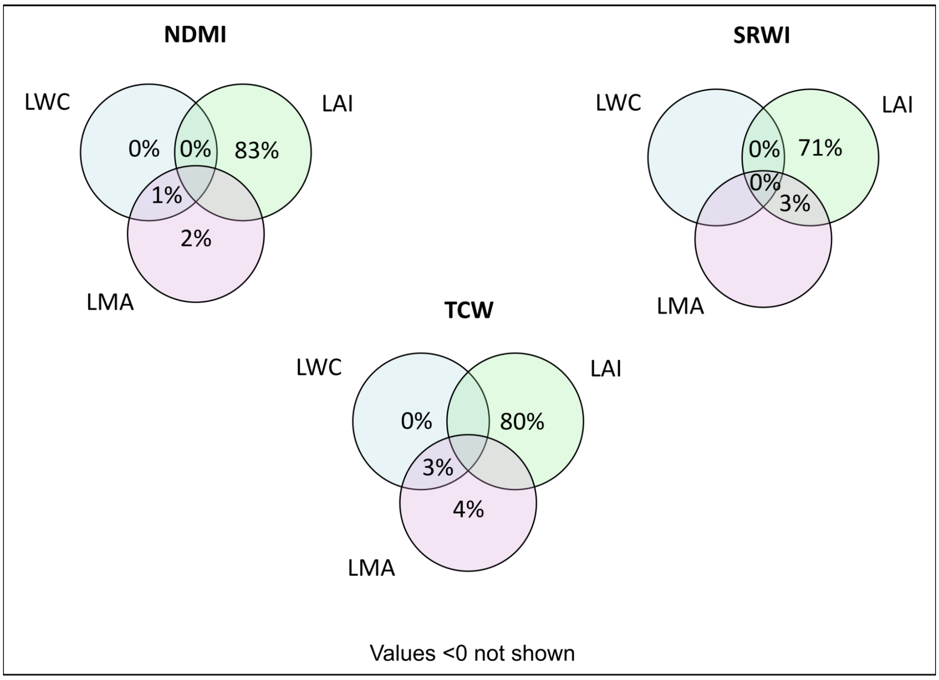


**Figure S4.** Variance partitioning analysis for three water-related multispectral indices—NDMI, SRWI and TCW—based on ground-truth and satellite data. Each circle represents a predictor variable included in the models: leaf water content (LWC), leaf area index (LAI), and leaf mass per area (LMA). Numbers indicate the percentage of variance in each index uniquely or jointly explained by these traits. The high independent contribution of LAI across indices indicates that variation in water-related indices is predominantly influenced by canopy structure rather than by leaf water content.

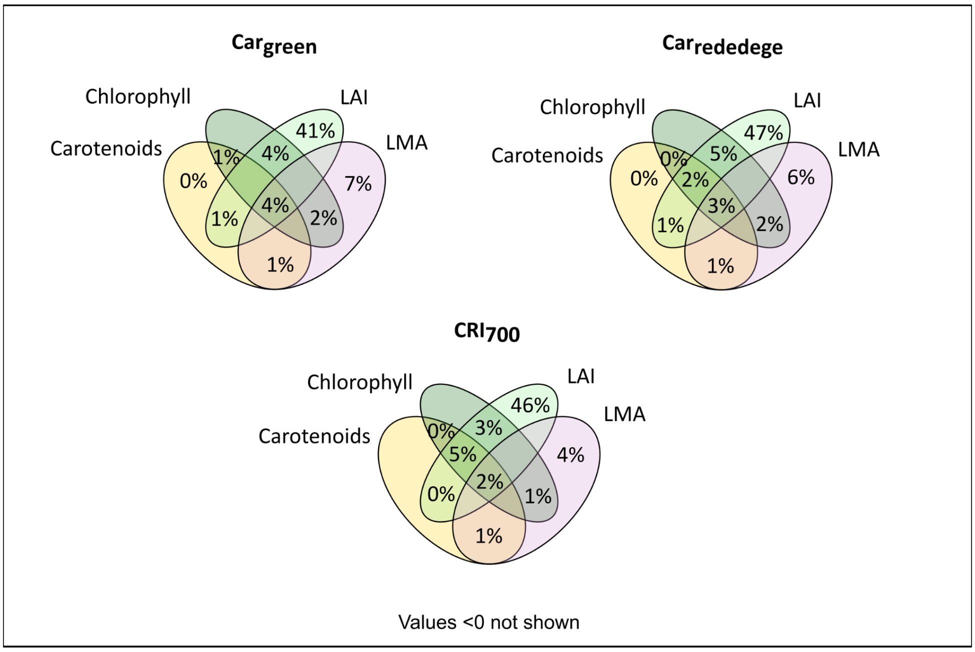


**Figure S5.** Variance partitioning analysis for three carotenoid-related multispectral indices — $Car_{green}$, $Car_{rededge}$, and $CRI_{700}$ — based on ground-truth and satellite data. Each circle represents a predictor variable included in the models: leaf carotenoid content, leaf chlorophyll content, leaf area index (LAI), and leaf mass per area (LMA). Numbers indicate the percentage of variance in each index uniquely or jointly explained by these traits. The large independent contribution of LAI and the shared effects with chlorophyll indicate that variation in carotenoid-related indices is strongly influenced by canopy structure and a small fraction of pigment co-variation rather than by carotenoid content alone.

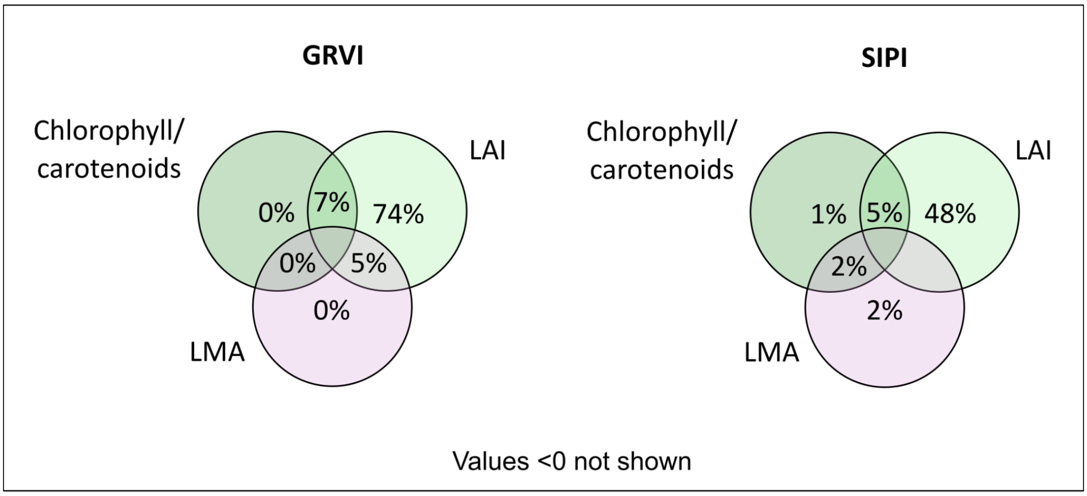


**Figure S6.** Variance partitioning analysis for two multispectral indices associated with the chlorophyll/carotenoid ratio — SIPI and GRVI — based on ground-truth and satellite data. Each circle represents a predictor variable included in the models: leaf chlorophyll/carotenoid ratio, leaf area index (LAI) and leaf mass per area (LMA). Numbers indicate the percentage of variance in each index uniquely or jointly explained by these traits. The strong independent contribution of LAI and the low fractions explained by pigments indicate that variation in these indices is largely driven by canopy structure rather than by the chlorophyll/carotenoid ratio.

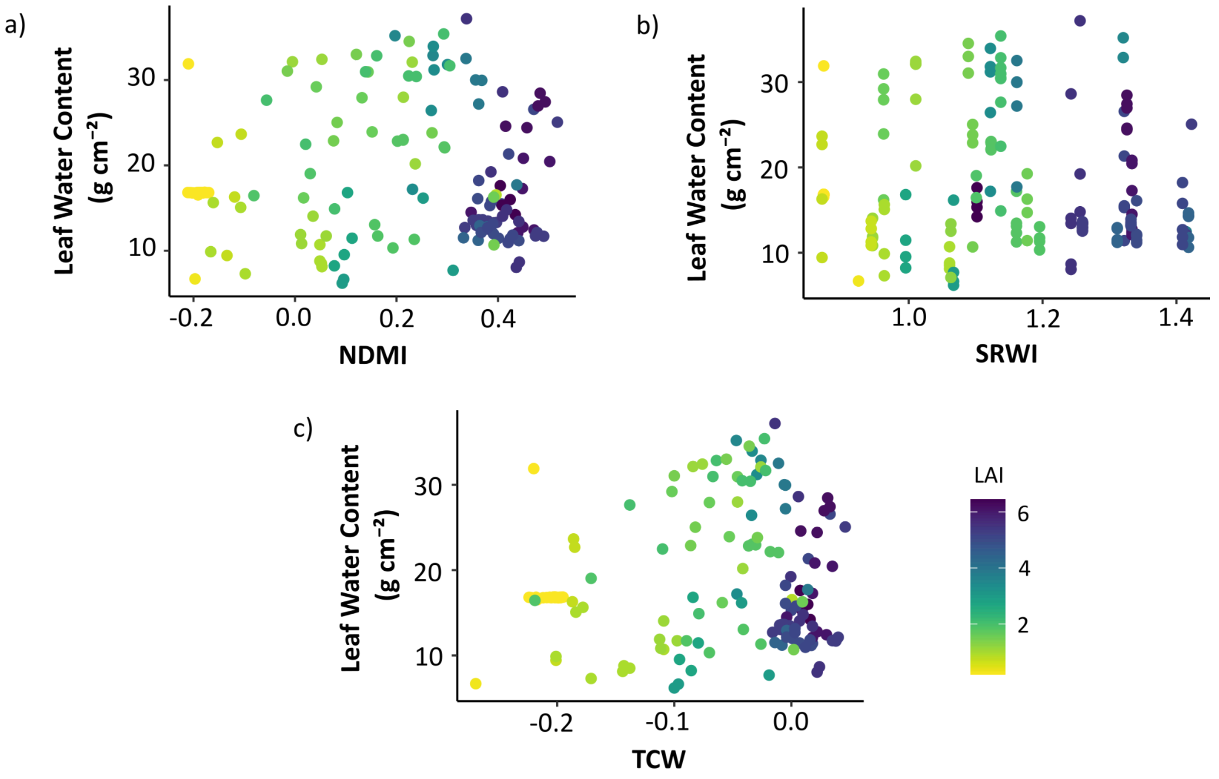


**Figure S7.** Relationships between three water-related multispectral indices—SRWI, NDMI, and TCW—and leaf water content, colored by leaf area index (LAI). Each point represents a field–satellite paired observation. These indices were evaluated under the same conditions as those in the main results but are presented here for completeness.

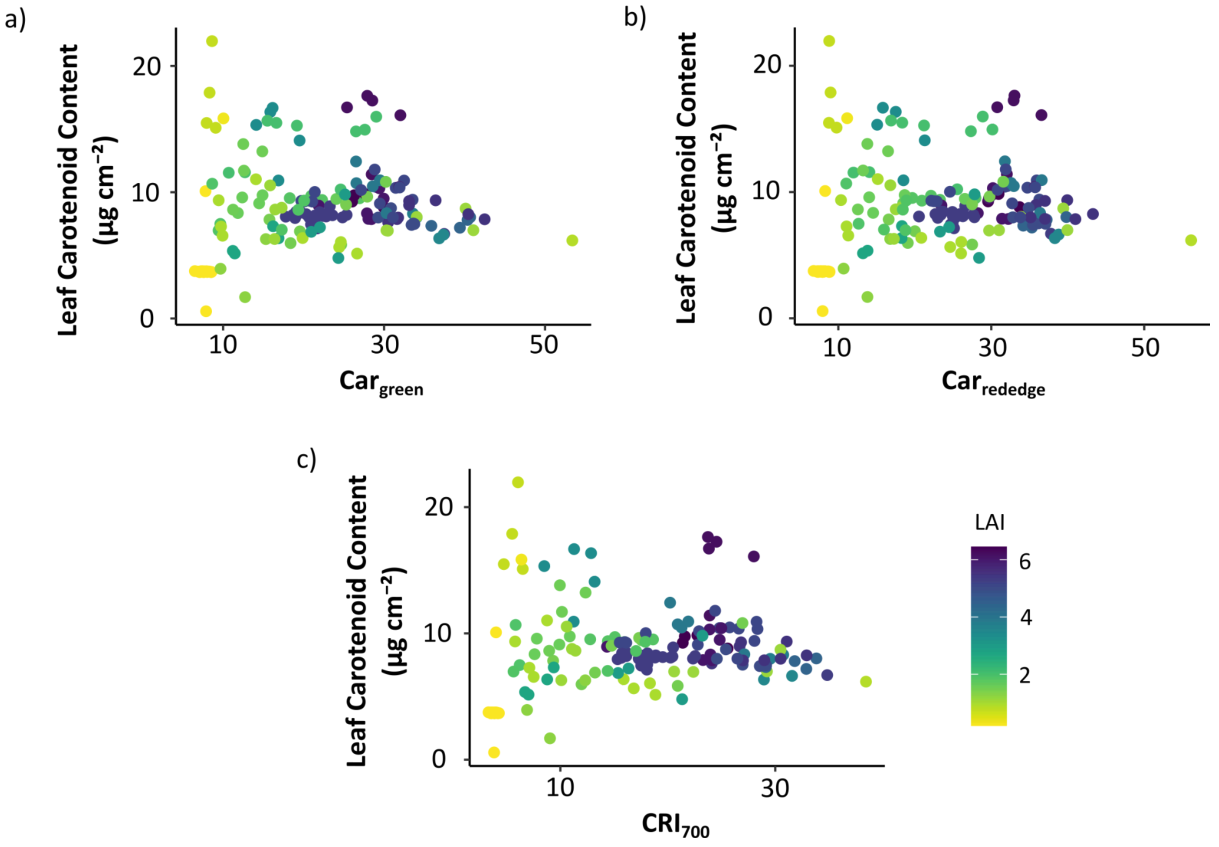


**Figure S8.** Relationships between three carotenoid-related multispectral indices—$CRI_{700}$, $Car_{green}$, and $Car_{rededge}$—and leaf carotenoid content, colored by leaf area index (LAI). Each point represents a field–satellite paired observation. These indices were evaluated under the same conditions as those presented in the main results, but are shown here for completeness.

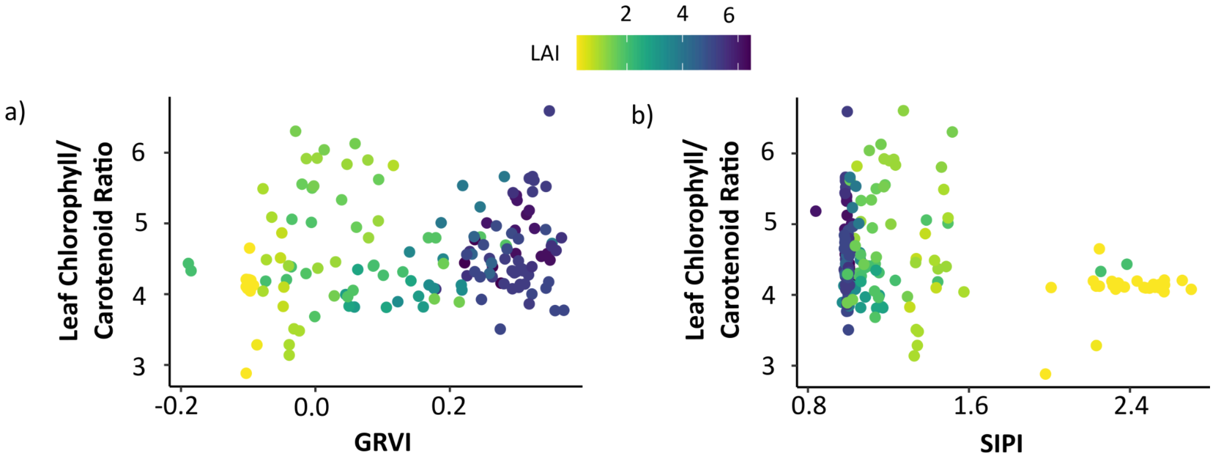


**Figure S9.** Relationships between two multispectral indices associated with the chlorophyll/carotenoid ratio—SIPI and GRVI—and the leaf chlorophyll/carotenoid ratio, colored by leaf area index (LAI). Each point represents a field–satellite paired observation. These indices were evaluated under the same conditions as those in the main results but are shown here for completeness.
